# Uncovering structural determinants of peptide recognition by public and private T-cell receptors

**DOI:** 10.1101/2025.11.23.690020

**Authors:** Caner Akıl, Yanchun Peng, Julia McCarthy, Yanan Zhu, Nathan Hardenbrook, Danning Dong, Elie Antoun, Xuan Yao, Guihai Liu, Ji-li Chen, Esra Balıkçı, Raul Cioaca, Yunkai Yang, Alexander J. Mentzer, Julian C. Knight, Ricardo Fernandes, Tao Dong, Peijun Zhang

**Author notes:** These authors contributed equally (Caner Akıl, Yanchun Peng, Julia McCarthy, Yanan Zhu).

## Abstract

Public T cell receptors (TCRs) recurrently emerge across individuals in response to common pathogens, yet the structural and biophysical basis distinguishing public from private clonotypes remains incompletely defined. Here, we combine epitope mapping, single-cell TCR sequencing, and single-particle cryo-electron microscopy to dissect CD8^+^ T cell responses to the immunodominant SARS-CoV-2 ORF3a(207–215) epitope presented by HLA-A*01:01. Among responding clonotypes, we identify a shared public TCR (TCRpub) and an individual-specific private TCR (TCRpriv) that use nearly identical TRBV5-1 β chains but distinct α chains. Both clonotypes exhibit comparable micromolar affinity and functional avidity, yet their structures reveal different antigen-recognition modes. We determined cryo-EM structures of the TCR_pub_ and TCR_priv_ in complex with ORF3a_(207-215)_/ HLA-A*01:01 at ∼3 Å resolution. Despite targeting the same epitope, the two receptors engaged the peptide-MHC complex with distinct CDR-loop orientations and contact footprints: TCRpub engages the peptide through a peptide-centric AGDL CDR3β motif and focuses interactions on the MHC α2-helix, whereas TCRpriv distributes contacts across both MHC α-helices via a canonical CDR3β configuration. These findings illustrate how near-identical β chains can yield divergent recognition strategies to recognise the same pMHC ligand through alternative α-chain pairing. More broadly, this work establishes cryo-EM as a robust approach for resolving physiological-affinity TCR/pMHC complexes, providing mechanistic insight into how public TCRs emerge and persist in antiviral immunity.

## INTRODUCTION

T cells are central to adaptive immunity, detecting infected or malignant cells through TCRs that engage peptide-MHC (pMHC) complexes on target cells^1–6^. Somatic recombination of variable (V), diversity (D), and joining (J) gene segments generates vast receptor diversity, which is refined by thymic selection to yield a peripheral repertoire of ∼10^8^ clonotypes capable of broad pathogen recognition while maintaining self-tolerance^7–11^. Despite this immense diversity, TCRs with identical or highly similar sequences, termed public TCRs, often recur across unrelated individuals responding to the same antigen. Such convergence likely reflects recombination biases and selection pressures that favour particular αβ pairings with optimal structural complementarity to specific pMHCs, leading to advantageous functional outcomes. Strikingly, public TCR responses are frequently observed against immunodominant epitopes from common viral infections, including SARS-CoV-2^12–20^. Previous studies, including our own^21^, have shown that public TCRs are associated with high-frequency, pre-existing precursors^15,22^ and may contribute to rapid and broad early immune responses^14,15^. Public clonotypes have also been implicated in autoimmune diseases^23,24^, but remain notably rare in cancer, despite extensive repertoire profiling efforts. In contrast, private TCRs, which are unique to individuals, dominate most immune responses. However, the structural and functional distinctions between public and private clonotypes, and how these differences influence recognition of identical peptide-MHC ligands, remain poorly defined^25^.

Although thousands of TCR sequences with known antigen specificity have been catalogued, structural coverage remains sparse and biased. Nevertheless, key principles of TCR/pMHC engagement have emerged: hypervariable CDR3 loops primarily contact the bound peptide, whereas germline-encoded CDR1 and CDR2 loops interact mainly with the MHC helices^26–32^. Most TCR/pMHC complexes have been solved by X-ray crystallography, a method that favours high-affinity or engineered receptors and relies on extensive recombinant protein production. This has introduced a notable skew toward certain V-gene usage, leaving much of the natural sequence space unexplored^33,34^. Consequently, many physiologically relevant TCRs, particularly those with low-to-moderate affinities, remain structurally uncharacterized due to challenges in obtaining crystals of sufficient quality. Single-particle cryo-electron microscopy (cryo-EM) now provides a powerful complement to crystallography, enabling visualisation of macromolecular complexes in solution. Recent advances have extended the limits of cryo-EM, enabling near-atomic resolution of smaller assemblies and opening new avenues for studying TCR/pMHC interactions across a broader affinity spectrum.

To explore how closely related clonotypes can adopt distinct antigen-recognition modes, we examined two CD8^+^ T cell receptors from convalescent SARS-CoV-2 patients that recognise the same ORF3a_(207–215)_ epitope (FTSDYYQLY), presented by the common HLA-A*01:01 allele. Recognition of this epitope elicits strong and frequent CD8^+^ T cell responses in convalescent individuals, with a high degree of clonotype sharing across donors. Within this response, we identified two dominant clonotypes, one public (TCR_pub_; TRAV3/TRBV5-1) and one private (TCR_priv_; TRAV25/TRBV5-1), that recognise the same pMHC ligand yet differ only in their α-chain usage and four CDR3β residues. This natural pair provides an ideal system to dissect how α-chain pairing and subtle CDR3 differences shape antigen recognition within a shared β-chain scaffold.

Here, we combine epitope mapping, single-cell TCR sequencing, affinity measurements, and high-resolution cryo-EM to resolve the structures of TCR_pub_ and TCR_priv_ bound to ORF3a_(207–215)_/HLA-A*01:01. Despite comparable micromolar affinities and similar overall docking architectures, the two receptors engage the peptide-MHC surface through strikingly divergent CDR-loop configurations. Together, our findings establish cryo-EM as a robust approach to visualise low-affinity, wild-type TCR/pMHC complexes and uncover how alternative α-chain pairings reconfigure the antigen-binding site, and shape antigen recognition by the β-chain.

## RESULTS

### Dominant SARS-CoV-2 ORF3a_(207-215)_-A*01:01-specific T cells utilise a high frequency of public TCRs

Previous studies conducted by us and others have identified a dominant CD8^+^ epitope ORF3a_(207-215)_ (FTSDYYQLY) derived from the SARS-CoV-2 ORF3a protein restricted by HLA-A*01:01^39–42^. In our cohort, 52 individuals who recovered from COVID-19 were HLA typed, and 18 (34.6%) were HLA-A*01:01 positive. An *ex vivo* interferon (IFN)-γ ELISpot assay showed that 88.9% (16/18) of these individuals responded to this epitope, representing 30.8% (20/52) of the overall cohort (**Fig. 1a, Supplementary Table 1)**. To characterise this dominant T cell response, ORF3a_(207-215)_-A*01:01-specific T cells from four donors were sorted at the single-cell level by flow cytometry using peptide-MHC Pentamer for Smartseq2 sequencing (**Supplementary Fig. 1a and 1b**). Analysis of their TCR usage revealed a high frequency of shared TCR chains detected in more than one individual (Supplementary Data 1). As shown in **Fig. 1b-c**, 26 out of 85 TRA (30.6%) chains and 6 out of 46 β chains (13%) were shared across multiple individuals. When considering full αβ pairings, only a subset of these constituted public TCR clonotypes (shared complete αβ TCRs).

**Figure 1.**
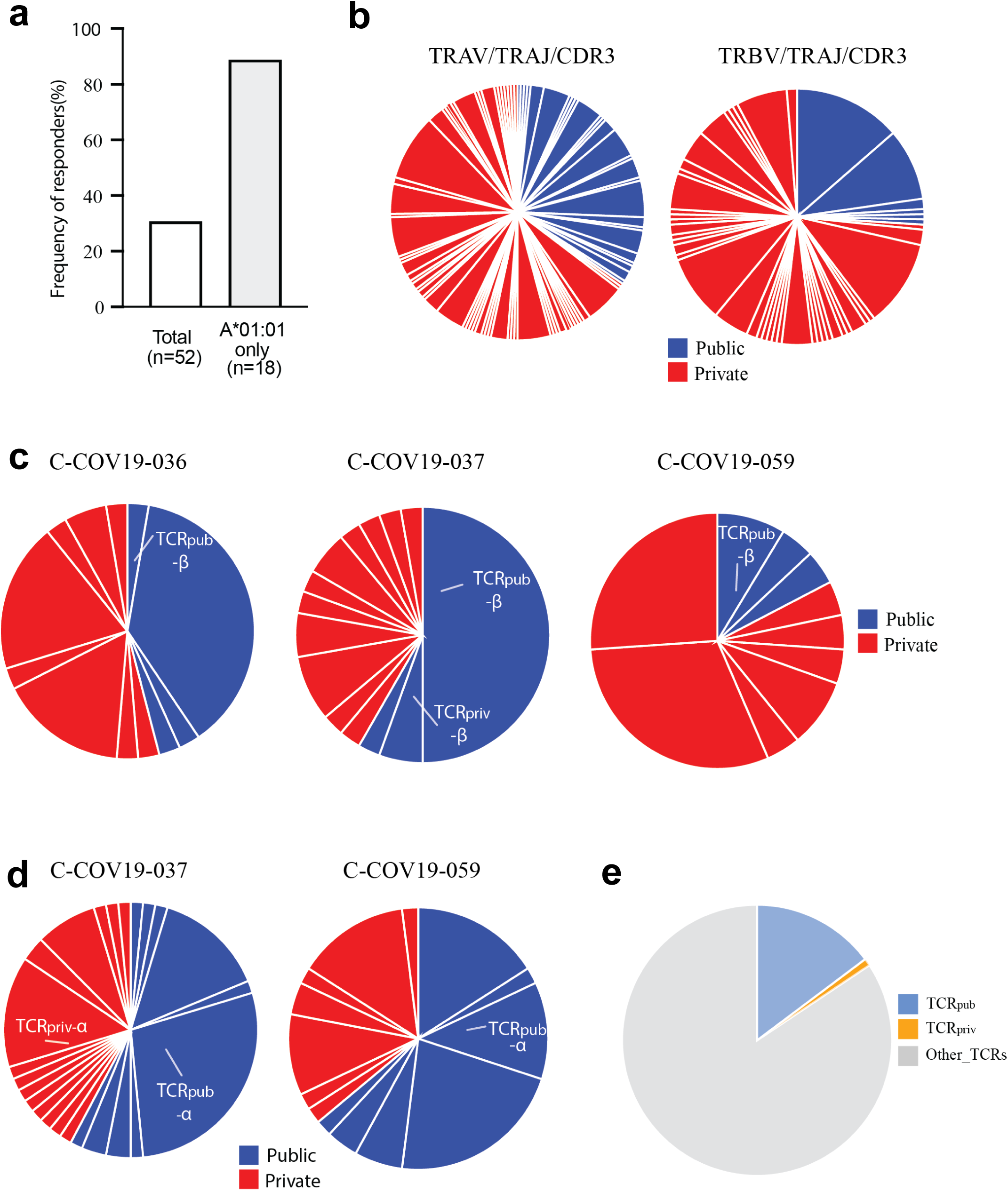
Identification of a public TCR specific to ORF3a_(207-215)_/HLA-A*01:01. (**a**) Frequency of responders to the HLA-A01:01-restricted ORF3a_(207-215)_ epitope (FTSDYYQLY). (**b**) Frequency of public T cell receptors by TRAV/TRAJ/CDR3A (left) and TRBV/TRBJ/CDR3B (right) usage in ORF3a_(207-215)_/HLA-A01:01-specific CD8⁺ T cells. (**c**) Frequencies of TCR_pub_ β chain and TCR_priv_ β chain in three individuals. (**d**) Frequencies of TCR_priv_ and TCR_pub_ α chains in four individuals. (**e**) Frequencies of paired TCR_priv_ and TCR_pub_ in ORF3a_(207-215)_/HLA-A*01:01-specific cells across four individuals. TCR_priv_ = 1, TCR_pub_ = 17, other TCRs = 94.

Among these TCRs, two TCRs exhibit particularly interesting characteristics (**Supplementary Table 2**). Both TCRs utilise a similar β chain (TRBV5-1/TRBJ1-1) with slightly different CDR3 variable regions. One of the TCRs is termed as public T cell receptor (TCR_pub_) as the paired αβ chains were detected in two individuals. TCR_pub_ contains a shared β chain that was detected in three individuals: C-COV19-036 with a frequency of 2.70%, C-COV19-037 with a frequency of 50%, and C-COV19-059 with a frequency of 8.7%), whereas another TCR (termed TCR_priv_) contains a unique β chain, only detected in one donor C-COV19-037 at 5.56% (**Fig. 1c**). TCR_priv_ employs a unique α chain (TRAV25/TRAJ57) detected only in C-COV19-037, whereas TCR_pub_ uses a shared α chain (TRAV3/TRAJ6) that detected in both C-COV19-037 and C-COV19-059 (**Fig. 1d**). Interestingly, TCR_pub_ was detected at a higher frequency (17.5%) than the TCR_priv_ (1%) (**Fig. 1e**).

We also analyzed the COVID-19 OTS dataset, and this analysis revealed that 148 and 92 TCRs shared CDR3β sequences similar to TCR_priv_ and TCR_pub_, respectively (Levenshtein similarity score > 0.8). The β chains in these TCRs were paired with 244 and 146 unique α chains, respectively, indicating a greater α-chain diversity among TCR_priv_-like TCRs. Conversely, 94 and 45 TCRs exhibited CDR3α similarity to TCR_priv_ and TCR_pub_, pairing with 196 and 83 unique β chains, respectively. The maximum cross-chain similarity scores were 0.733 (α) and 0.692 (β) for TCR_priv_, and 0.714 (α) and 0.769 (β) for TCR_pub_, suggesting that while none of the pairings reached high absolute similarity, distinct diversity patterns emerged between the two receptors. Overall, TCR_priv_-like repertoires displayed broader combinatorial diversity with highly variable α-β pairings. In contrast, TCR_pub_-like repertoires were characterized by more recurrent and conserved α-β combinations, consistent with their classification as private and public TCRs, respectively (**Fig. 2a, b**).

**Figure 2.**
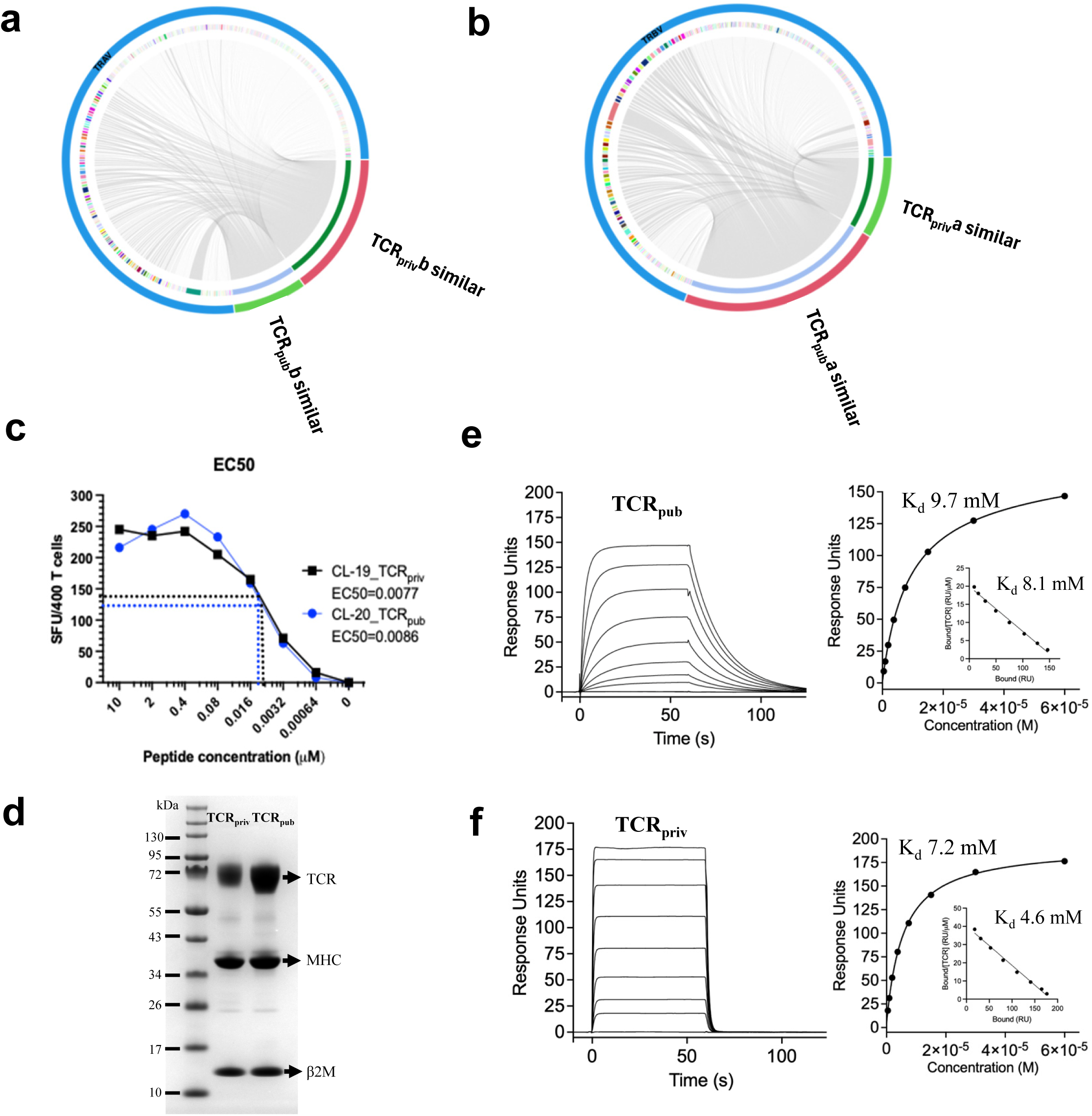
Biochemical characterization of TCR/pMHC complexes and TCR repertoire similarity in COVID OTS dataset. (**a-b**) CDR3 sequence similarity and α-β pairing diversity among OTS COVID TCRs. (**a**) Circos plot showing TRAV gene usage and α-chain pairing of OTS COVID TCRs that share similar β CDR3 sequences with TCR_priv_ and TCR_pub_ (Levenshtein similarity > 0.8). TCRs with CDR3β similar to TCR_priv_ (148 sequences) and TCR_pub_ (92 sequences) are indicated as TCR_priv_b_similar (pink) and TCR_pub_b_similar (green), respectively. Their paired α chains are connected by grey links, illustrating highly diverse α chain usage (244 and 146 unique α CDR3s, respectively). (**b**) Circos plot showing TRBV gene usage and β-chain pairing of OTS COVID TCRs that share similar α CDR3 sequences with TCR_priv_ and TCR_pub_. TCRs with α CDR3 similar to TCR_priv_ (94 sequences) and TCR_pub_ (45 sequences) are shown as TCR_priv_a_similar (pink) and TCR_pub_a_similar (green). Grey links connect these α-similar TCRs to their paired β chains (196 and 83 unique CDR3βs, respectively), revealing a more conserved α-β pairing pattern in TCR_pub_-like repertoires. (**c**) T cell clones expressing TCR_pub_ and TCR_priv_ show similar EC50 values. (**d**) SDS–PAGE gel (4–12% Bis-Tris gradient gel, run with MES buffer) showing purified TCR_pub_/pMHC and TCR_priv_/pMHC complexes under reducing conditions. (**e-f**) Left: SPR sensorgrams showing binding of eight concentrations of TCR_pub_ (**e**) and TCR_priv_ (**f**) (starting at 60 μM with 2-fold serial dilutions) to 220-250 RU of immobilized ORF3a₍₂₀₇₋₂₁₅₎–HLA-A*01:01 complexes at 37 °C. Responses from a reference flow cell containing an irrelevant pMHC were subtracted from the total signals. Right: Nonlinear curve fitting of the untransformed data using a 1:1 Langmuir binding model yielded a Kd = 7.2 μM and Bmax = 176 RU for TCR_pub_ (**e**), and Kd = 9.7 μM and Bmax = 147 RU for TCR_priv_, respectively. A linear Scatchard plot (inset) produced a similar Kd = 4.6 μM (**e**) and Kd = 8.1 μM (**f**). Data are representative of three independent experiments.

### TCR_pub_ and TCR_priv_ recognize ORF3a_(207-215)_-HLA-A*01:01 with micromolar affinity and comparable functional avidity

To assess the functional sensitivity of TCR_pub_ - and TCR_priv_-expressing T cell clones, we evaluated IFN-γ responses using an ELISpot assay. ORF3a**_(207-215)_-**A*01:01 specific T cell clones were isolated by single-cell with peptide-MHC multimer sorting, *in vitro* expansion and validated by multimer staining. Clones with >95% purity were used for experiments (**Supplementary Fig. 1b and 1c**). Clone 19 (TCR_priv_) and Clone 20 (TCR_pub_) exhibited similar functional avidities, with EC_50_ values of 0.0077 μM and 0.0086 μM, respectively (**Fig. 2c, Supplementary Fig. 1d**). To dissect how TCR_pub_ and TCR_priv_ clonotypes engage ORF3a_(207-215)_-A*01:01 (hereafter referred to as the peptide-MHC complex, pMHC), we generated soluble TCR heterodimers by removing the transmembrane regions and co-expressing paired α- and β-chains in mammalian cells (**Supplementary Fig. 2a**). These soluble receptors enabled both biophysical and functional interrogation of ORF3a-specific TCR/pMHC interactions. Two representative complexes, TCR_pub_/pMHC and TCR_priv_/pMHC were used for further detailed characterization (**Fig. 2d**, **Supplementary Fig. 2b-d**). Sequence alignments confirmed that, as expected from their shared TRBV gene usage, the TCR β-chains are highly conserved between these clonotypes, with differences limited to the four amino acids in the CDRβ3 loop. In contrast, the TCR α-chains exhibit greater overall sequence diversity (**Supplementary Fig. 3a,b**).

Binding affinities of TCR_pub_ and TCR_priv_ to the same cognate pMHC complex were quantified by surface plasmon resonance (SPR), revealing moderate affinities in the range of 7.2 – 9.7 µM (**Fig. 2e, f**). To further characterize the biochemical properties of the soluble, recombinant TCRs, mass photometry analysis of TCR_pub_/pMHC and TCR_priv_/pMHC complexes revealed that both complexes are predominantly monomeric with a small fraction (∼5%) of TCR_pub_/pMHC complex exhibiting a dimer configuration (**Supplementary Fig. 2e, f**).

### Cryo-EM structures of TCR_pub_ and TCR_priv_ in complex with ORF3a_(207-215)_-HLA-A*01:01

We performed single-particle cryo-EM analysis of TCR_pub_/pMHC, TCR_priv_/pMHC complexes specific to the ORF3a_(207-215)_ epitope. To address the challenges associated with structural determination of these natural TCR/pMHC complexes, including their small size (∼93 kDa), inherent flexibility, low affinity, preferred orientation, and particle disassembly at the air-water interface, we collected a large dataset for each complex (4-6 million particles) and additionally recorded images at a 30° tilt angle. To minimize particle heterogeneity and orientation bias, we further carried out iterative rounds of particle picking, followed by stringent two-dimensional (2D) and three-dimensional (3D) classifications (**Supplementary Fig. 4 and 5, Supplementary Table 3)**. Final reconstructions produced cryo-EM maps of TCR_pub_/pMHC and TCR_priv_/pMHC complexes, with global resolutions of 3.17 Å and 3 Å, respectively **(Supplementary Fig. 4, 5**). The resulting cryo-EM maps of TCR_pub_/pMHC and TCR_priv_/pMHC exhibited well-resolved features for the TCR α and β chains, the ORF3a_(207-215)_ peptide and the α1/α2 helices of the MHC (**Fig. 3**). The MHC α-chain and β-microglobulin of TCR_pub_/pMHC complex were not well-resolved, likely due to their structural flexibility in the TCR_pub_/pMHC complex. In both TCR_pub_/pMHC and TCR_priv_/pMHC complexes, the ORF3a_(207-215)_peptide is buried in the cleft formed by the MHC α1 and α2 helices. The peptide is mainly contacted by the CDR3α and CDR3β loops **(Fig. 4b, e)**, while the MHC is primarily engaged by CDR1 and CDR2 loops of both TCRα and TCRβ chains **(Fig. 4c, f)**.

**Figure 3.**
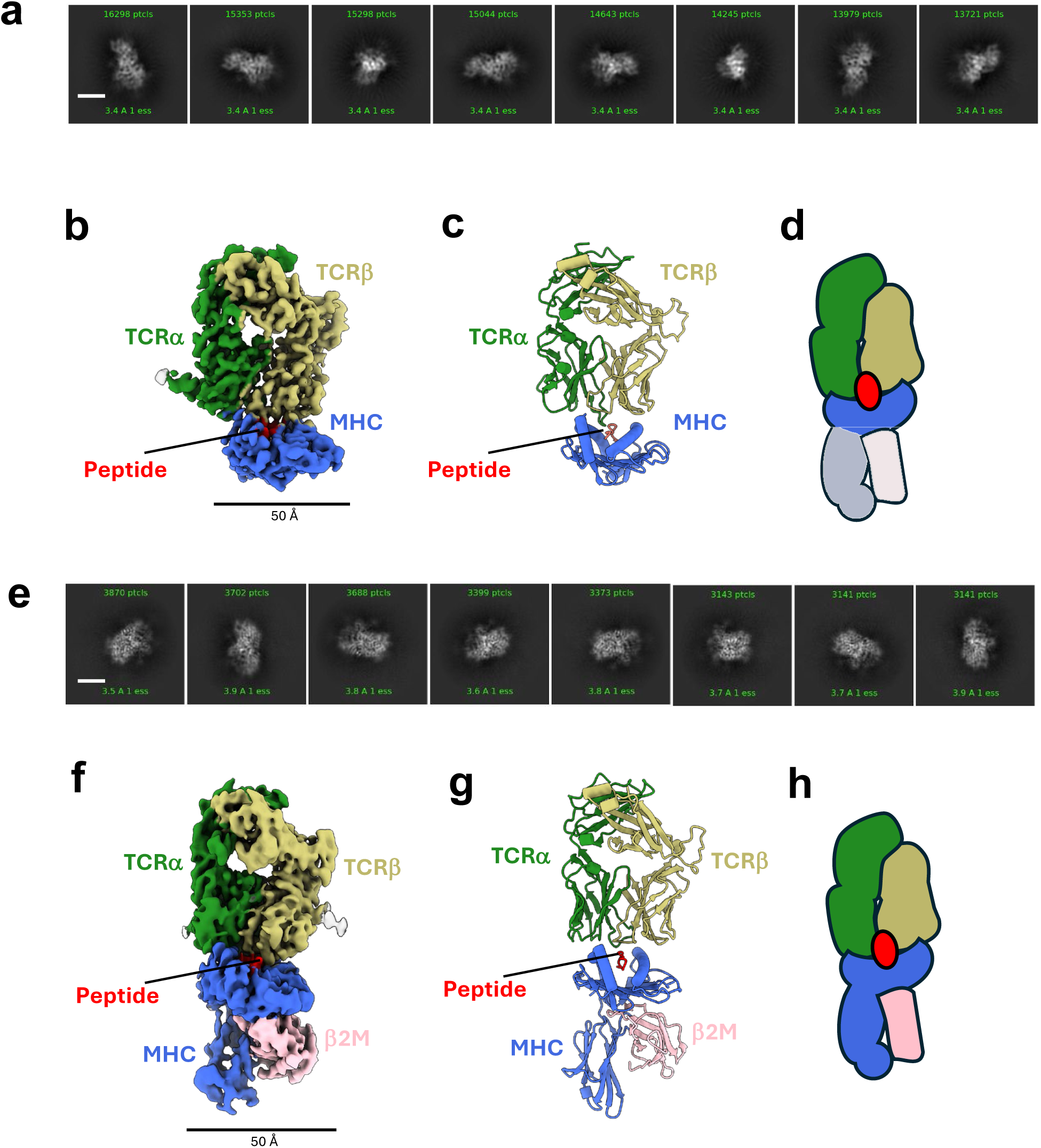
Cryo-EM structures of TCR_pub_/pMHC and TCR_priv_/pMHC complexes. (**a**) Representative 2D class averages of TCR_pub_/pMHC. (**b**) Cryo-EM density map of TCR_pub_/pMHC with subunits shown in indicated colors. (**c**) Ribbon diagram of the TCR_pub_/pMHC structural model. (**d**) Cartoon schematic of TCR_pub_/pMHC. The TCRα chain is depicted in green, the TCRβ chain in dark khaki, the peptide ligand in red, and the MHC molecule in blue. The pMHC (dark grey) and β2-microglobulin (light grey) were not well resolved, likely due to their flexibility. **(e)** Representative 2D class averages of TCR_priv_/pMHC. (**f**) Cryo-EM density map of TCR_priv_/pMHC with subunits shown in indicated colors. (**g**) Ribbon diagram of the TCR_priv_/pMHC structural model. (**h**) Simplified cartoon schematic of TCR_priv_/pMHC. Scale bars in a and e (white), and in b and f (black) indicate 50 Å.

**Figure 4.**
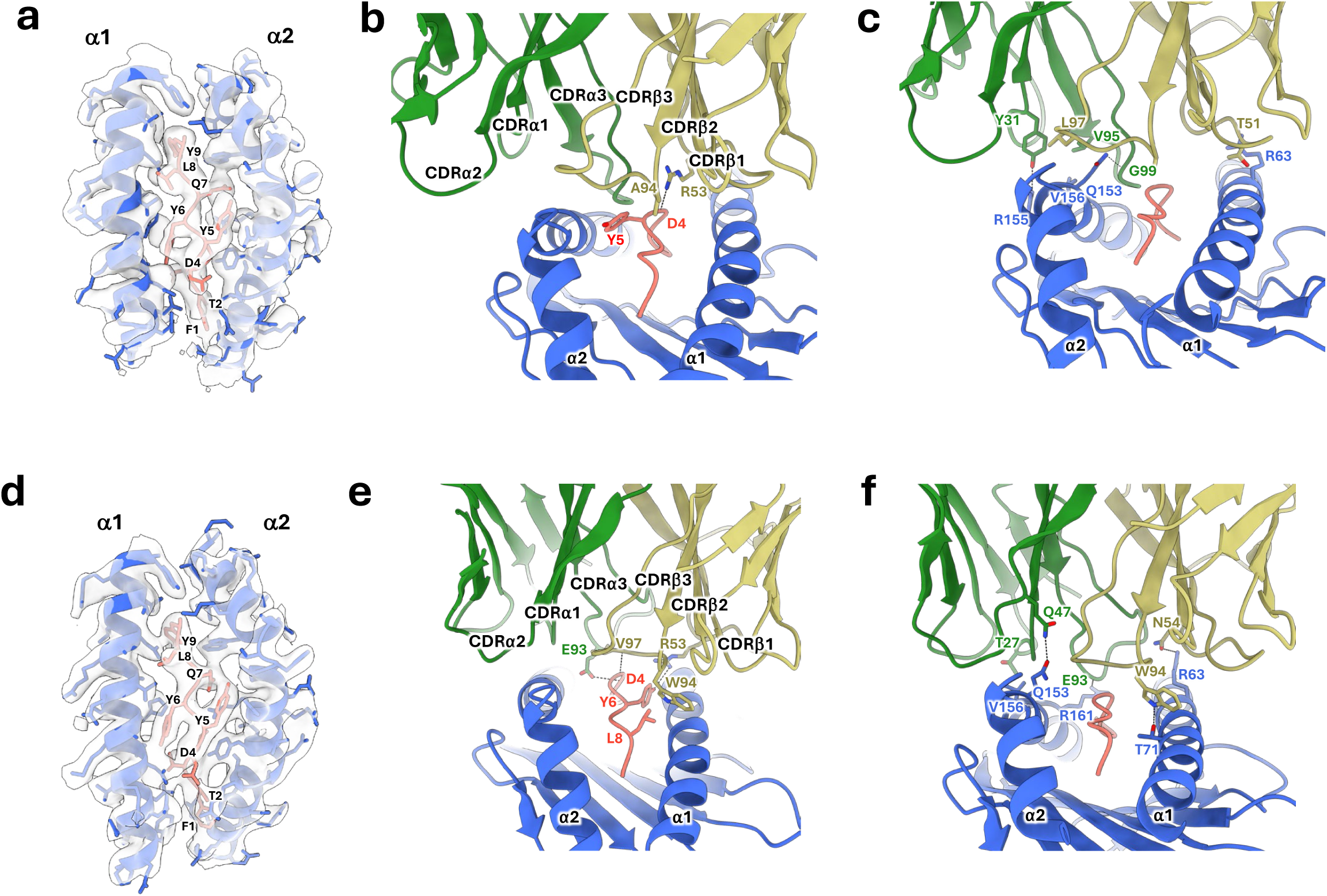
TCR recognition of the ORF3a_(207-215)_ peptide and MHC molecule. (**a-c**) Structural details of the TCR_pub_/pMHC complex with density and model superimposed, shown at the ORF3a_(207-215)_ peptide (FTSDYYQLY) and MHC interface (**a**), the interface between TCR_pub_ CDR loops and the peptide with interacting residues labelled (**b**), and the interface between TCR_pub_ CDR loops and MHC with interacting residues labelled (**c**). **(d-f)** Structural details of the TCR_priv_/pMHC complex with density and model superimposed, shown at the peptide and MHC interface (**d**), the interface between TCR_priv_ CDR loops and the peptide with interacting residues labelled (**e**), and the interface between TCR_priv_ CDR loops and MHC with interacting residues labelled (**f**). The TCRα chain is shown in green, the TCRβ chain in dark khaki, the peptide ligand in red, and the MHC molecule in blue. The TCRα chain CDR loops (CDRα1, CDRα2, and CDRα3) and TCRβ chain CDR loops (CDRβ1, CDRβ2, and CDRβ3) are marked.

**Figure 5.**
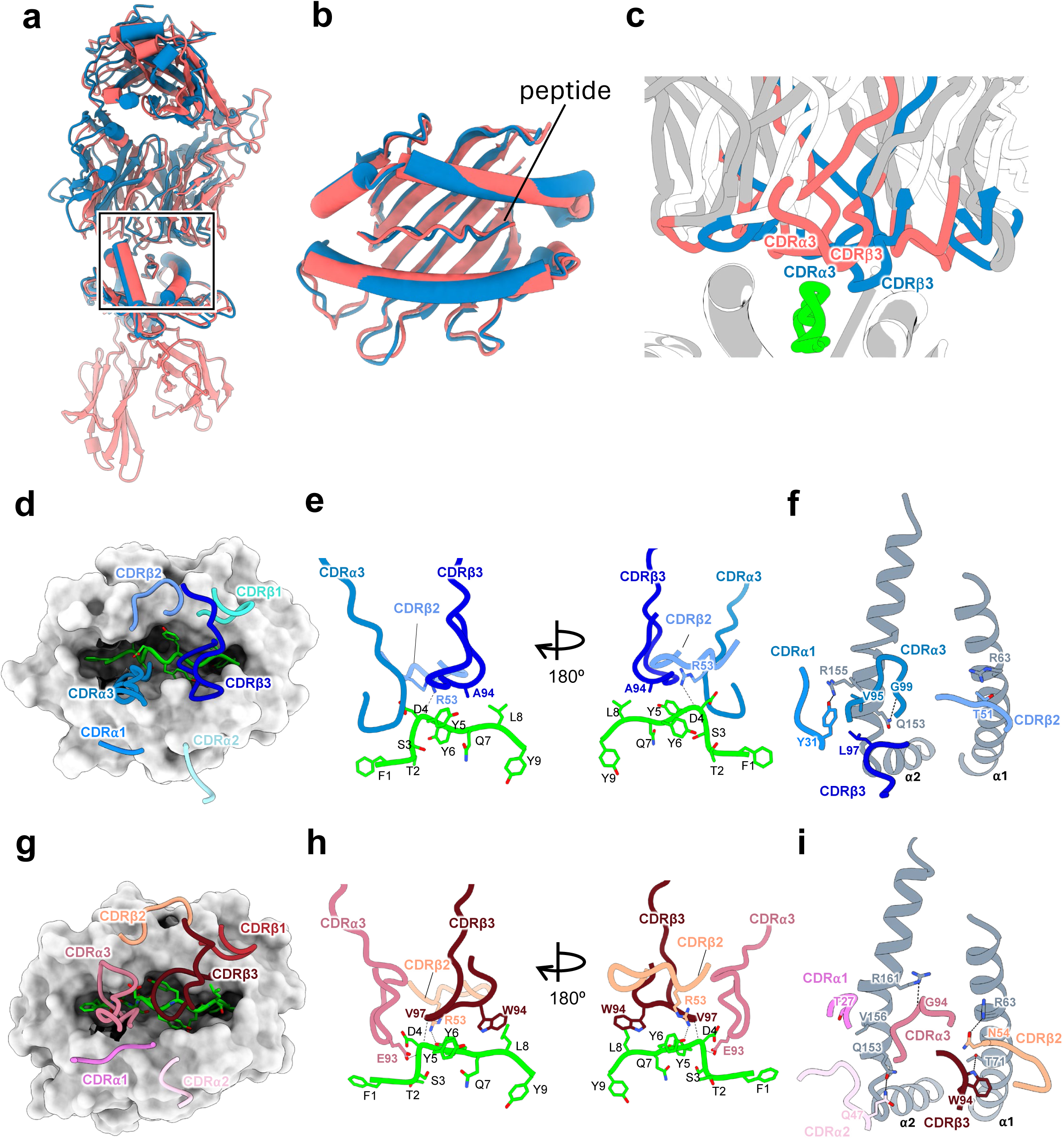
Comparative analysis of TCR_pub_ and TCR_priv_ engaging peptide antigen and MHC. (**a**) Superimposition of TCR_pub_/pMHC (blue) and TCR_priv_/pMHC (salmon) and complexes aligned on the pMHC. (**b**) Zoom-in on the aligned pMHC region. (**c**) Zoom-in on the CDR loops from TCR_pub_/ (blue) and TCR_priv_ (salmon) . The peptide is shown in green. (**d**) Footprint of TCR_pub_/ CDR loops (blue tones) on the pMHC complex (grey surface). (**e-f**) Interaction between the TCR_pub_ CDR loops (blue tones) and the peptide (green, **e**) and the MHC (grey, **f**). Hydrogen bonds are shown with dashed lines. (**g**) Footprint of TCR_priv_ CDR loops (red tones) on the pMHC complex (grey surface). (**h-i**) Interaction between TCR_priv_ CDR loops and the peptide (green, **h**) and the MHC molecule (grey, **i**).

The local resolution at the peptide-binding cleft in both the TCR_pub_/pMHC and TCR_priv_/pMHC complexes was sufficiently high to allow confident modelling of side chains (**Fig. 4, Supplementary Fig. 6)**, enabling detailed analysis of the molecular interactions of these wild-type TCR/pMHC complexes that underpin T cell recognition. The conformations of the MHC, the ORF3a_(207-215)_peptide, and their interfaces were nearly identical in both complexes (**Fig. 4a, d**). The N-terminus of the peptide forms hydrogen bonds with Tyr residues on the MHC, while Asp4, Tyr5, and Tyr6 engage the MHC helices, all contribute to stabilizing the peptide-MHC interaction **(Fig. 4a, d, Supplementary Fig. 6)**.

### Modes of antigen recognition by TCR_pub_ and TCR_priv_

Detailed interface analysis revealed that both TCR_pub_ and TCR_priv_ recognise antigen through a distinct combination of peptide- and MHC-mediated contacts. In the TCR_pub_/pMHC complex, the backbone of the peptide residue Asp4 forms a hydrogen bond with Arg53 in the CDRβ2 loop of the TCR. Additionally, Ala94 in the CDRβ3 loop makes a moderate hydrophobic interaction with Tyr5 of the peptide (**Fig. 4b**). For TCR_pub_ and MHC interactions, Val156 in the MHC α2 helix is positioned to form a hydrophobic contact with Val95 in the CDRα3 loop, while Thr51 in the CDRβ2 loop engages in a hydrophobic interaction with Arg63 in the MHC α1 helix. Additionally, Gln153 and Arg155 in the MHC α2 helix form hydrogen bonds with Gly99 in the CDRα3 loop and Tyr31 in the CDRα1 loop, respectively (**Fig. 4c**).

In the TCR_priv_/pMHC complex, the peptide residue Tyr6 forms a hydrogen bond with Arg53 in the CDRβ2 loop. In this conformation, the backbone of Asp4 also potentially forms a salt bridge with the side chain of Arg53, stabilizing the central portion of the interface (**Fig. 4e**). Additional backbone-backbone hydrogen bonds are observed between Asp4 and Val97 in the CDRβ3 loop, and Glu93 in the CDRα3 loop (**Fig. 4e**). Furthermore, Leu8 of the peptide engages Trp94 in the CDRβ3 loop through hydrophobic interactions. The TCR CDR loops engage the antigen peptide more extensively in TCR_priv_ than in TCR_pub_. For TCR_priv_/MHC interactions, multiple hydrogen bonds are observed between the TCR and both α1 and α2 helices of the MHC. Gln47 in the CDRα2 loop forms a hydrogen bond with Gln153 of the MHC α2 helix; Asn54 in the CDRβ2 loop forms a salt bridge with Arg63 of the α1 helix; the backbone of Glu93 forms a hydrogen bond with Arg161 of the α2 helix; Trp94 in the CDRβ3 loop contacts Thr71 of the α2 helix; and Thr27 in the CDRα1 loop engages Val156 through a hydrophobic interaction (**Fig. 4f**). Notably, Asp4 of the peptide is a shared contact residue in both TCR_pub_ and TCR_priv_, underscoring its central role in mediating TCR-peptide recognition. In contrast, Tyr5 is uniquely contacted by TCR_pub_, while Tyr6 and Leu8 are specific to TCR_priv_. Collectively, these contacts stabilize the TCR/pMHC interface and contribute to antigen specificity.

### Public and private TCRs exhibit divergent solutions to antigen recognition despite nearly identical β chains

While the overall architectures of the TCR_pub_/pMHC and TCR_priv_/pMHC complexes are nearly the same, with an RMSD of 1.13 Å (**Fig. 5a, b**), their CDR interaction profiles differ markedly (**Fig. 5c**). Importantly, the CDRβ3 loop of TCR_pub_ is positioned much closer to the peptide than that of TCR_priv_, potentially contributing to its distinct peptide recognition (**Fig. 5c**), whereas CDRs of TCR_priv_ establish more extensive contacts with the MHC α-helices. This differential interaction pattern may underlie the flexibility and versatility of the TCR_pub_/pMHC complex.

To delineate the interaction profiles of individual CDR loops, we projected the TCR-peptide-MHC interfaces from a top-down perspective and mapped the contact footprints over the pMHC surface. Although both complexes adopt a similar overall docking configuration, distinct differences were observed in the contributions of individual CDR loops to the peptide and MHC recognition. Notably, the CDRα3 and CDRβ3 loops exhibited divergent engagement patterns (**Fig. 5d, g**). In TCR_pub_/pMHC, the CDRβ3 loop adopts a distinct AGDL motif (**Fig. 5e, Supplementary Fig. 3a**), replacing the WTGV motif found in TCR_priv_ (**Fig. 5h**). A key difference lies in Trp94 of the WTGV motif in TCR_priv_, which mediates dual contacts with Leu8 of the peptide and Thr71 of the MHC (**Fig. 5h-i**). This residue is replaced by Ala in TCR_pub_, eliminating these interactions (**Fig. 5e-f**). However, the altered positioning of TCR_pub_ allows a hydrogen bond to form between the backbone atoms of Arg53 (CDRβ2) and Asp4 of the peptide (**Fig. 5e**). Additionally, the AGDL motif supports hydrophobic interactions between Ala94 (CDRβ3) and Tyr5 of the peptide (**Fig. 5e**), resulting in a more peptide-centric docking for the TCR_pub_. On the other hand, Arg53 in the CDRβ2 loop of TCR_priv_ interacts with Tyr6 of the peptide (**Fig. 5e**), a contact absent in TCR_pub_ (**Fig. 5e**). Interestingly, TCR_pub_ interactions are primarily concentrated on the α2 helix of the MHC (**Fig. 5f**), whereas TCR_priv_ engages both the α1 and α2 helices more evenly (**Fig. 5i**). This difference may reflect the greater structural flexibility of the TCR_pub_ conformation. Collectively, these findings establish a structural framework by which distinct CDR loop engagements shape differential peptide and MHC recognition, as well as the overall flexibility of TCR_pub_ versus TCR_priv_ complexes.

To assess the structural variability of antigen recognition, we compared the TCR_pub_/pMHC and TCR_priv_/pMHC complexes against a representative set of class I TCR/pMHC structures from TCR3d database^43^, selected on the basis of TM score^44^ and restricted to HLA-A alleles to maintain the MHC background. The TM-score, which ranges from 0 to 1 (with values above 0.5 generally indicating similar overall folds), is a length-independent metric that evaluates global topological similarity between two protein structures and is less sensitive to local deviations compared with RMSD (see Methods). TCR_priv_/pMHC exhibited relatively greater structural similarity to previously characterised complexes, as reflected by lower RMSD values, consistent with a more canonical binding mode. Conversely, TCR_pub_/pMHC showed higher RMSD values relative to the reference set, indicating a more divergent docking geometry. Overall, TCR_pub_/pMHC preserved the overall architecture of class I TCR/pMHC complexes but displayed greater local variability at the TCR/pMHC interface and within the pMHC α1 and α2 domains (**Fig. 6**). Data from this specific pair of TCR_pub_/pMHC and TCR_priv_/pMHC complexes suggest that public TCRs may adopt distinct CDR engagement modes that optimize structural complementarity and confer selective advantages during repertoire formation.

**Figure 6.**
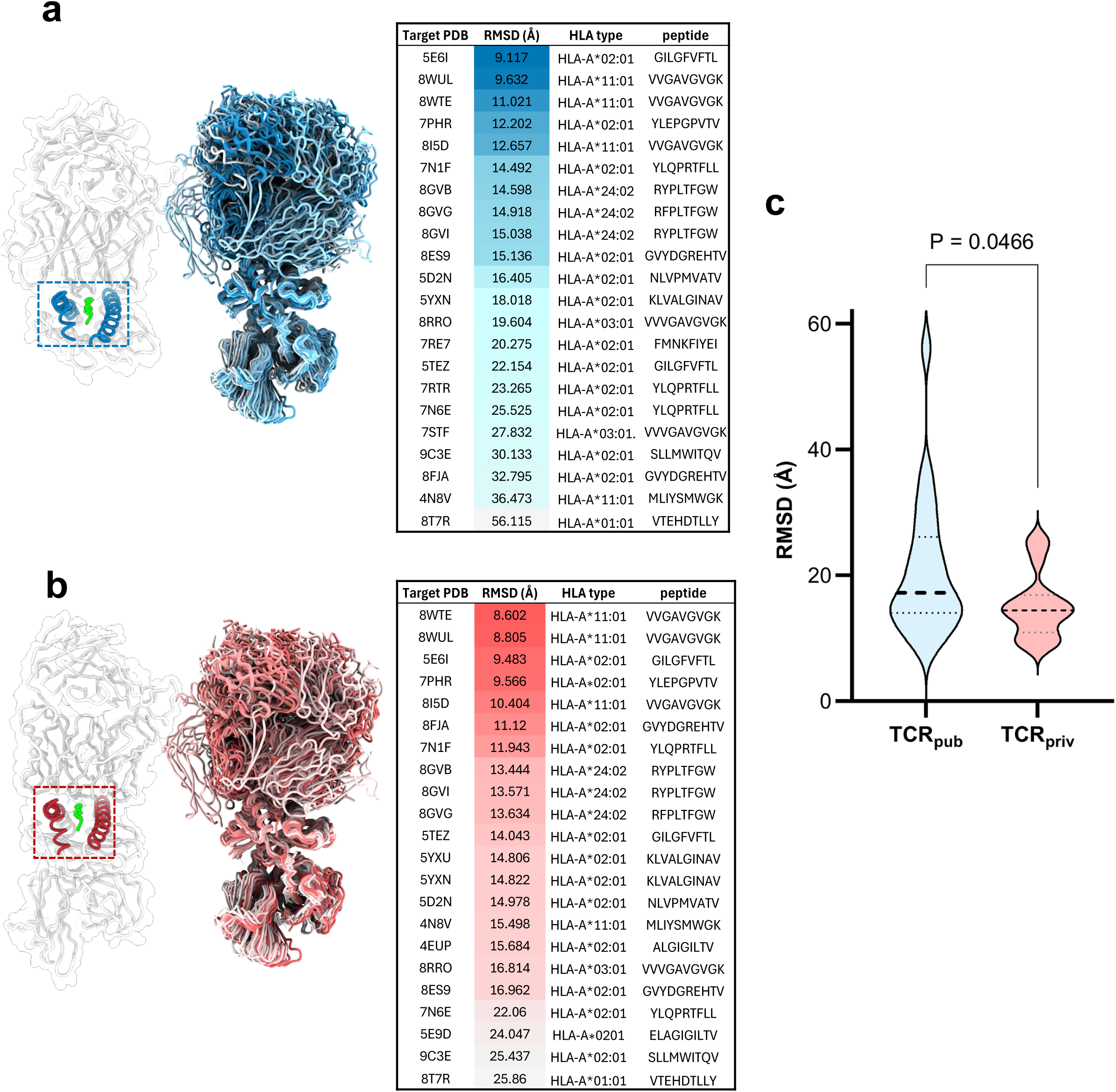
Structural flexibility of TCR/pMHC complexes. (**a**) Structural alignment of the TCR_pub_/pMHC complex against 1,388 class I TCR/pMHC structures from the TCR3d database. TM-align was used to identify structures with the highest topological similarity based on TM-scores. The top hits were superimposed on the MHC α1 and α2 helices (highlighted in red boxes, left panel) and the bound peptide. RMSD values for the TCR α and β chains were calculated in ChimeraX. (**b**) The same strategy was applied to the TCR_priv_/pMHC complex to obtain RMSD values for its TCR α and β chains. (**c**) Comparison of RMSD values for TCR_pub_/pMHC (n = 50) and TCR_pub_/pMHC based on the reference structures. Data are shown as medians. A two-tailed Mann–Whitney test indicated a significant difference between the two groups (U = 157, p = 0.0466; *p < 0.05). Median RMSD values: for TCR_pub_/pMHC (n = 50) and TCR_pub_/pMHC= 17.21, TCR_priv_/pMHC = 14.42. Hodges–Lehmann difference: –3.693 (95.07% CI: –8.287 to –0.0660).

## DISCUSSION

Understanding how TCRs decode pMHCs is central to explaining the balance between diversity and reproducibility in antiviral immunity^3^. Although public TCRs are repeatedly observed across unrelated individuals, the structural logic that enables these clonotypes to emerge with such convergence remains unclear. By resolving near-native structures of two wild-type TCRs recognizing the same SARS-CoV-2 ORF3a_(207–215)_/HLA-A*01:01 complex, we reveal how closely related clonotypes, sharing an almost identical β chain, can nevertheless adopt distinct docking solutions that differentially partition peptide and MHC contacts. These findings provide an example for how conserved β-chain scaffolds can support multiple α-chain-dependent recognition modes, offering new insight into the emergence of public TCRs.

A central observation from our work is that TCRpub engages the pMHC through a peptide-focused recognition strategy, whereas TCR_priv_ distributes its footprint more broadly across both MHC α-helices via a canonical binding interface. These distinct solutions arise despite similar overall docking geometry (RMSD 1.13 Å) and comparable micromolar affinities, demonstrating that public versus private behaviour cannot be inferred from affinity alone. Instead, α-chain pairing emerges as the key determinant: the TRAV3 α chain in TCR_pub_ rotates and reorients the CDR3α–CDR3β interface, relocating the β-chain loops to favour peptide-centric binding. By contrast, the TRAV25 α chain in TCR_priv_ preserves a more canonical architecture that stabilizes balanced MHC contacts.

More broadly, this work demonstrates that cryo-EM can resolve wild-type, physiological-affinity TCR/pMHC complexes without the need for affinity enhancement, stabilization, or supraphysiological engineering. X-ray crystallography has long been used to provide atomic views of TCR/pMHC interfaces, but this has also resulted in a limited and biased dataset, reflecting the challenges posed by the need to obtain diffracting crystals and the production of significant amounts of TCR/pMHCs^45^, which can obscure key aspects of cross-reactivity^46^, specificity, and immune escape^47^. Using large-particle datasets and tilt-collection strategies, we achieved 3.0–3.2 Å Cryo-EM reconstructions that captured native CDR-loop conformations and fine-scale differences at the peptide interface. Recent advances in cryo-EM have resolved full-length TCRαβ-CD3, TCR/pMHC-co-receptor, and TCRγδ-CD3 structures^48–51^. However, such studies often depend on affinity-enhanced TCRs or Fab fragments, limiting physilogical wild-type TCR/pMHC interactions. In the context of viral immunity, such limitations may overlook physiological responses to subdominant or variable epitopes^52^. Bridging this gap requires structural information on wild-type TCR/pMHC complexes exhibiting physilogical affinity. These advances open the door to systematically mapping the structural landscape of bona fide antiviral TCRs, including those with affinities traditionally considered “too weak” for structural study. Such access is essential for understanding cross-reactivity, precursor selection, immune escape, and the conformational plasticity that often defines antiviral recognition.

Here, we report cryo-EM structures of two wild-type sequence TCRs in complex with the SARS-CoV-2 ORF3a_(207-215)_ epitope presented by HLA*01:01. These structures revealed how nearly identical TRBV5-1 β chains pair with distinct α chains (TRAV25 in TCR_priv_, TRAV3 in TCR_pub_), resulting in private and public TCR clonotypes. Both receptors engage the same peptide-MHC ligand with comparable functional avidity and micromolar binding affinity, demonstrating that different clonotypic architectures can mediate equally effective antiviral responses. Clonotype-specific CDR loop rearrangements further shape distinct binding geometries, illustrating how TCR diversity accommodates a single antigen through multiple docking modes and thereby expands the functional repertoire of the T cell response.

The recurrence of TCR_pub_ in multiple donors classifies it as a public clonotype: TRBV5-1 CDR3β motifs pair with diverse α chains in various contexts, whereas the TRAV25 chain of TCR_priv_ remains uniquely linked to its β partner. By contrast, TRAV3-like α chains are broadly distributed across many β chains. These patterns indicate convergent antigen-driven selection and frequent recombination of public TCR motifs within the repertoire. Consistent with these observations, analysis of the COVID-19 OTS dataset revealed distinct repertoire features between TCR_priv_ and TCR_pub_. Although all α-β similarity scores remained below the 0.8 threshold, TCR_priv_-associated sequences exhibited broader combinatorial diversity, consistent with a private, individual-specific response. In contrast, TCR_pub_-associated sequences displayed more conserved α-β pairings, indicative of a shared or public clonotype recurring across multiple individuals. It should be noted that the 0.8 threshold is an operational cutoff reflecting ∼2 amino acid differences, and even “private” TCRs may occasionally appear in multiple individuals when large population datasets are examined. These observations suggest that repertoire architecture and pairing diversity, rather than absolute sequence similarity, are key determinants distinguishing private and public TCR responses in COVID-19 (Fig 2a, b).

Similar patterns, where a dominant α chain pairs with multiple diverse β chains encompassing both public and private clonotypes, were also observed in our recent study^21^. Although these analyses were based on bulk β-chain sequencing and paired naive datasets remain limited, such findings suggest that certain public motifs pre-exist in the naive pool and are preferentially expanded upon antigen exposure^21,54^. Future studies combining single-cell paired repertoire sequencing with structural and functional analyses will be valuable to clarify how public and private TCRs are generated, selected, and maintained across individuals.

This study exemplifies cryo-EM as an effective alternative to X-ray crystallography for natural TCR/pMHC recognition at micromolar affinity. We anticipate that the cryo-EM approach will be broadly adoped for more native and physiologically relevant TCR/pMHC complexes. Extending this approach to additional public and private TCR/pMHC pairs across diverse HLA alleles will systematically map the TCR conformational landscape. Integrating high-resolution structures with deep sequencing repertoires and biophysical profiling promises predictive models of TCR specificity and cross-reactivity, potentially informing next-generation adoptive T cell therapies.

## METHODS

### Study participants and ethics

Patients were recruited from the John Radcliffe Hospital in Oxford, UK, between March 2020 and September 2021 by identification of patients hospitalized during the SARS-CoV-2 pandemic and recruited into the Sepsis Immunomics study. Blood samples were collected at least 28 days after symptom onset during the primary infection as described previously^15^. Written informed consent was obtained from all patients. Ethical approval was given by the South Central-Oxford C Research Ethics Committee in England (ref. 19/SC/0296).

### IFN-γ ELISpot assay

*Ex vivo* IFN-γ ELISpot assays were performed using either freshly isolated or cryopreserved PBMCs, or antigen-specific T cell clones, as previously described^15^. For *ex vivo* assays, peptides (Genscript, UK) were added at a concentration of 2 μg ml^-1^ to 2 × 10□ PBMCs per well and incubated for 16-18 hours. Negative control wells were set up without peptide stimulation. For assays using T cell clones, autologous EBV-transformed B cell lines (BCLs) were pre-pulsed with peptides at five fold titrated concentrations and subsequently co-cultured with T cells at an effector-to-target (E: T) ratio of 1:50 for at least 6 hours. Co-cultures of EBV-transformed B cells and T cell clones without peptide stimulation were included as negative controls. Antigen-specific responses were quantified using AID ELISpot 7.0 software. The mean spot number from negative control wells was subtracted from peptide-stimulated wells, and results were expressed as spot-forming units (s.f.u.) per 10 PBMCs for ex vivo ELISpot or s.f.u. per 400 T cells when using T cell clones. Responses were considered positive if the spot count was at least three times the mean of the negative control wells. Assays were excluded from further analysis if background responses in negative control wells exceeded 30 s.f.u. per 10^6^ PBMCs or per 400 T cells, or if positive control wells did not yield a measurable response.

### Flow cytometric sorting of ORF3a_(207-215)_ -A*01:01-specific CD8^+^ T cells

ORF3a_(207-215)_-HLA-A01:01-specific CD8^+^ T cells were stained with a phycoerythrin (PE)-conjugated HLA-A*01:01 ORF3a_(207-215)_ pentamer (ProImmune). Nonviable cells were excluded using Live/Dead Fixable Aqua dye (Invitrogen). After washing, cells were stained with the following surface antibodies: CD3-FITC (BD Biosciences), CD8-PerCP-Cy5.5, CD14-BV510, CD19-BV510, and CD16-BV510 (BioLegend). Following exclusion of nonviable and CD14^+^, CD16^+^, and CD19^+^ cells, CD3^+^ CD8^+^ pentamer cells were sorted directly into 96-well PCR plates (Thermo Fisher Scientific) using a BD FACS Fusion or BD FACSAria III sorter (BD Biosciences) and stored at −80 °C for subsequent analyses.

### SmartSeq2 scRNA-seq and data processing

ScRNA-seq with *ex vivo* sorted CD8^+^ pentamer^+^ T cells (Proimmune, UK) was performed using SmartSeq2 ^55^ with the following modifications: reverse-transcription (RT) and PCR amplification were performed as previously described^55^with the exception of using ISPCR primer with biotin tagged at the 5′ end and increasing the number of cycles to 25. Sequencing libraries were prepared using the Nextera XT Library Preparation Kit (Illumina) and sequencing was performed on Illumina NextSeq sequencing platform with NextSeq Control Software v.4. BCL files were converted to FASTQ format using bcl2fastq v.2.20.0.422 (Illumina). FASTQ files were aligned to human genome hg19 using STAR v.2.6.1d^56–58^.

### SmartSeq2 TCR repertoire analysis

TCR sequences were reconstructed from scRNA-seq FASTQ files using MiXCR v.3.0.13 ^59,60^ generating separate TRA and TRB output files for downstream analysis. These output files were imported into R using the tcR package v.2.3.2. For paired αβ TCRs, cells were filtered to retain either one α and one β chain (1α1β) or two α chains and one β chain (2α1β). Additionally, lists of all cells expressing one β chain (regardless of the number of α chains) were generated for downstream analysis.

### Deep sequencing of TCR repertoire of T cell clones

T cell clones were generated by sorting CD8^+^ pentamer^+^ T cells into 96-well Tissue culture plates at single-cell level from thawed PBMCs. T cell clones were then expanded and maintained with irradiated allogeneic PBMCs every 2-3 weeks as described previously^61^. Purity and antigen-specificity of T cell clones was checked by staining with peptide-MHC tetramers. T cell clones were included in the experiments only if their purity exceeded 95%. From each T cell clone, 1 × 10^5^ cells were harvested and washed with phosphate-buffered saline (PBS). Total RNA was extracted using RNeasy Plus Micro Kit (QIAGEN), and 100 ng of total RNA from each T cell clone was used to generate full-length TCR repertoire libraries for Illumina Sequencing using a SMARTer Human TCR a/b Profiling Kit (Takara) following the supplier’s instructions. The cDNA sequences corresponding to variable regions of TCRα and/or TCRβ transcripts were amplified with primers, including Illumina indices, allowing for sample barcoding. PCR products were then purified using AMPure beads (Beckman Coulter). The quantity and quality of cDNA libraries were checked on an Agilent 2100 Bioanalyzer system. Sequencing was performed using MiSeq reagent Kit v.3 (600 cycles) on MiSeq (Illumina) with MiSeq Control Software v.2.6.2.1.

BCL files were converted to FASTQ files as described earlier. TCRs were extracted using MiXCR and the resulting output files (TRA and TRB) were parsed into R using tcR as described earlier. TCRs were filtered to retain 1α1β for each clone^62^.

### Synthesis of peptide-MHC complex

4 mg of the peptide FTSDYYQLY (synthesized by Genscript, 90% purity) was used. The HLA-A*01:01 heavy chain cDNA was modified by replacing the transmembrane and cytosolic regions with a sequence encoding the BirA biotinylation enzyme recognition site, as previously described ^63^. These modified heavy chains (amino acids 1-276) and β2-microglobulin were expressed in a prokaryotic system (pET; R&D Systems), purified from bacterial inclusion bodies, and solubilized in 8 M urea. Refolding was carried out by dilution in the presence of peptide at 4 °C for 48 hours. The refolded complexes were concentrated using an Amicon Stirred Cell with a 10 kDa ultrafiltration membrane (Millipore), then desalted on a PD-10 column (Cytiva) prior to biotinylation with BirA (Avidity). Monomeric complexes were further purified by gel filtration chromatography (Superdex 75, Cytiva), with the 45 kDa peak collected and concentrated to 1 mg/mL using an Amicon Ultra centrifugal filter (10 kDa cutoff, Millipore).

### Expression and purification of soluble TCRs

Synthetic dsDNA fragments encoding TCRα and TCRβ chains were obtained from Twist Bioscience, with overlaps designed for Gibson assembly into the pD649 vector (Addgene plasmid #156543). Vectors were linearised with restriction enzymes and assembled using Gibson Isothermal Assembly (NEB). Assembled plasmids were transformed into *E. coli* DH5α (NEB C2987H) and plated on LB agar containing 1% carbenicillin. Colonies were cultured overnight in LB medium, and plasmids were extracted and verified by Sanger sequencing (GENEWIZ). Recombinant TCRs were expressed in Expi293F cells (Thermo Fisher) maintained at 37°C with 5% CO_2_, shaking at 160 rpm. On the day of transfection, cells were diluted to 3.0 × 10 cells/mL in Expi293 medium (Thermo Fisher A1435101). Cells were transiently co-transfected with 25 µg of each TCRα and TCRβ plasmid using 150 µL of ExpiFectamine 293 reagent (Thermo Fisher), following the manufacturer’s protocol. At 18 h post-transfection, enhancers 1 and 2 were added. Supernatants were harvested on day 5, clarified by centrifugation (3,200g, 10 min, 4°C), and diluted 1:1 in HBS (30 mM HEPES pH 7.4, 150 mM NaCl). The buffer was supplemented with 20 mM Tris-HCl (pH 8.0) and 0.05% NaN . Proteins were captured by incubation with 3 mL of 50% Ni-NTA agarose (Qiagen 30210) overnight at 4°C with gentle agitation.

His-tagged TCRs were eluted by IMAC, concentrated to ∼500 µL, and treated overnight with HRV 3C protease (ab285999) to remove tags and leucine zippers. Proteins were further purified by size-exclusion chromatography (SEC) on a Superdex 200 Increase 10/300 GL column (Cytiva) using an ÄKTA Pure system equilibrated with HBS. Fractions were analysed by SDS-PAGE, and those with correct size and purity were pooled, concentrated (Amicon Ultra-4, 100 kDa MWCO, Sigma-Aldrich UFC8100), aliquoted, and flash-frozen in liquid nitrogen for storage at −80°C.

### Surface Plasmon Resonance

SPR measurements were carried out on a Biacore S200 system (Cytiva). Biotinylated ORF3a_(207-215)_-HLA-A*01:01 complex was immobilised captured on a CAP sensor chip using the Biotin CAPture Kit (Cytiva), yielding 150-250 RU. An irrelevant control protein was immobilised on a reference flow cell at a similar level. Serial twofold dilutions of soluble 3C-cleaved TCRs were injected at 20 μL/min in HBS-P+ running buffer (10 mM HEPES, 150 mM NaCl, 0.05% v/v Surfactant P20, pH 7.4) at 37°C) at 37°C. Binding responses at equilibrium were used to derive *K*_D_ values by fitting to a 1:1 Langmuir binding model using Biacore evaluation software. All measurements were performed in triplicate with alternating TCR injection order.

### Mass photometry analysis of TCR/pMHC complexes

Mass photometry measurements were performed at room temperature using a Refeyn TwoMP instrument (Refeyn Ltd, Oxford, UK). Prior to use, glass coverslips and silicone gaskets were thoroughly cleaned with HPLC-grade water followed by isopropanol, and dried under filtered air. TCR_priv_/pMHC and TCR_pub_/pMHC complexes were diluted to a final concentration of 200 nM in size-exclusion chromatography (SEC) buffer. To set the focal plane, 18 μl of buffer was first applied to the coverslip. Subsequently, 2 μl of the sample was added and data acquisition was initiated using medium camera image settings. Each acquisition consisted of a 1-minute movie, which was processed using ratiometric contrast analysis. Instrument calibration was carried out using reference standards including monomeric bovine serum albumin (66 kDa), dimeric BSA (132 kDa), and thyroglobulin (660 kDa), yielding a molecular weight accuracy within 5%. Experiments were independently repeated twice. Data were analyzed using DiscoverMP version 2.3 (Refeyn Ltd, Oxford, UK).

### Cryo-EM SPA sample preparation and data collection

Cryo-EM grids of TCR_priv_/pMHC and TCR_pub_/pMHC complexes were prepared by directly freezing the purified complexes at ∼0.5-1.0 mg/mL onto glow-discharged Quantifoil R1.2/1.3 Cu 300 mesh grids (Quantifoil Micro Tools GmbH) using a Vitrobot Mark IV (Thermo Fisher Scientific), without chemical crosslinking. Grids were blotted for 3.0 s at room temperature and 100% humidity, then plunge-frozen into liquid ethane. Grid quality was initially screened on a Thermo Scientific Glacios transmission electron microscope. Final data collection was performed on a Titan Krios transmission electron microscope (Thermo Fisher Scientific) operated at 300 kV. The TCR_pub_/pMHC complex was imaged using a Gatan K3 direct electron detector, while the TCR_priv_/pMHC complex were collected on a Falcon 4i direct electron detector equipped with a Selectris X energy filter in EER mode. The total electron dose was 50 e /Å², and images were collected with a defocus range of -0.8 to -2.5 μm. Micrographs were collected at a calibrated pixel size of 0.825 Å for the TCR_pub_/pMHC complex. In total, 12,657 and 10,222 movies were acquired for the TCR_pub_/pMHC (EMDB-54262, PDB 9RU5), TCR_priv_/pMHC (EMDB-54366, PDB 9RXM) (**Supplementary Table 3**).

### Cryo-EM single particle analysis

All datasets were processed using CryoSparc v4.4.1^64^, following standard single-particle analysis workflows, including motion correction, CTF estimation, iterative classification, and refinement. For the TCR_pub_/pMHC dataset, 12,657 micrographs were initially processed using patch motion correction and CTF estimation. Initial particle coordinates were generated via blob picking, and Bin2 particle extraction yielded 5,535,316 particles. After several rounds of 2D classification, 70,035 particles were retained. A Topaz model was then trained on these and used to reprocess the dataset. Following Bin2 re-extraction and additional 2D classification, 899,071 particles were obtained and refined down to 422,641 particles through further classification. These particles were subjected to ab initio reconstruction and heterogeneous refinement, resulting in two major classes containing 207,209 and 166,747 particles, respectively. The 207,209-particle class was selected for downstream analysis. Particles were re-extracted at Bin1, and focused 3D classification was performed. Classes displaying well-defined α-helices were selected for non-uniform refinement, followed by local refinement, yielding a global resolution of 3.17 Å at the FSC 0.143 criterion. Final post-processing included map sharpening and local resolution estimation (**Supplementary Fig. 4**).

For the TCR_priv_/pMHC dataset, a total of 10,222 movies were subjected to patch motion correction and CTF estimation. Initial particle coordinates were obtained using Topaz extraction with a model pre-trained on the TCR_pub_/pMHC dataset. Bin2 particle extraction yielded 3,903,441 particles, which were refined through multiple rounds of 2D classification, resulting in 89,350 particles for initial non-uniform refinement. To improve data quality, Topaz was retrained on high-quality particles from the initial rounds, and a new particle set was extracted and re-extracted at Bin1. Subsequent 2D classification using ViewSelect retained 457,735 particles. These were imported into RELION-5 for 3D auto-refinement with Blush regularization, yielding a global resolution of 3 Å at the FSC 0.143 criterion. Final post-processing involved map sharpening and local resolution estimation (**Supplementary Fig. 5**).

### Model building, refinement, and structure visualization

Initial atomic models of TCRα, TCRβ, the ORF3a_(207-215)_ peptide, and HLA-A*01:01 heavy chain were generated using AlphaFold3 multimer predictions^65^. For the TCR_priv_/pMHC and TCR_pub_/pMHC monomeric complexes, models were rigid-body docked into cryo-EM densities and refined iteratively using manual adjustments in Coot^66^ and real-space refinement in Phenix^67^. Structural figures were prepared using UCSF ChimeraX^68^.

### Sequence analysis

TCR_priv_ and TCR_pub_ α- and β-chain sequences were aligned using MUSCLE (v3.8)^69^ with default parameters. The alignment was visualized using ESPript 3.0^70^, with annotated CDR loops, and contact residues (**Supplementary Fig. 3a**). Structural comparisons between TCR_priv_/pMHC and TCR_pub_/pMHC were performed in ChimeraX (v1.7), and Cα RMSD values were calculated using the MatchMaker tool (**Supplementary Fig. 3b**). To assess structural similarity, TCR_priv_/pMHC and TCR_pub_/pMHC were aligned to 1,388 class I TCR-pMHC structures from TCR3d database^43^ using TM-align^44^. Alignments were performed by superposition on the MHC α1–α2 platform. To ensure comparable MHC backgrounds, only HLA-A–restricted complexes were retained for subsequent analysis, yielding a representative subset used for TM-score ranking and RMSD comparison (Supplementary Table 4). The best structural matches were selected based on TM-score and visual inspection, and pairwise RMSD values (Cα atoms) between TCR chains were computed in UCSF ChimeraX^68^ (Fig. 6). TM-scores (ranging from 0 to 1) were used to quantify global structural similarity independent of sequence length.

OTS COVID TCR sequence data were obtained from the Oxford Protein Informatics Group (OPIG) COVID-19 TCR database (https://opig.stats.ox.ac.uk/webapps/ots/ots_paired). The CDR3 sequences of TCR_priv_ α/β and TCR_pub_ α/β were pre-processed by removing the leading cysteine and terminal phenylalanine residues. Pairwise sequence similarity between each of the TCR_priv_ and TCR_pub_ α- and β-chain of CDR3 regions and all CDR3 sequences in the COVID OTS dataset was calculated using the stringsimmatrix function from the stringdist R package (v0.9.15), based on Levenshtein distance score. Similarity scores ranged from 0 (no similarity) to 1 (identical sequences). A threshold of >0.8, corresponding to approximately two amino acid differences, was applied to define sequences as “similar.” TCRs with CDR3α or CDR3β regions showing a similarity score >0.8 to TCR_priv_ or TCR_pub_ were classified as TCR_priv_ α/β-similar or TCR_pub_ α/β-similar, respectively. Paired α-β chain associations of these similar TCRs were visualized using Circos (v0.69) to illustrate V gene usage and pairing diversity. Distinct color segments represent V gene families (TRAV or TRBV), while grey connecting lines indicate α-β pairings (Fig. 2a-b; Supplementary Data 2).

### Quantification and statistical analysis

Statistical analyses were performed using GraphPad Prism (v10.3.0). EC_50_ values of T cell clones were calculated by nonlinear regression using a variable slope (four-parameter logistic) model in dose-response stimulation assays. Data are presented as mean ± SD or as medians, as indicated in the figure legends. Comparisons between two groups were performed using two-tailed Mann-Whitney tests. Effect sizes were estimated using the Hodges-Lehmann method with 95% confidence intervals. P values were corrected for multiple comparisons where applicable. Statistical significance is indicated as follows: *P ≤ 0.05, **P ≤ 0.005, ***P ≤ 0.0005, ****P ≤ 0.0001. Investigators were not blinded to group allocation during experiments or outcome assessment.

### Data availability

Cryo-EM maps and atomic models reported in this study have been deposited in the Electron Microscopy Data Bank (EMDB) and the RCSB Protein Data Bank (PDB). Accession codes and corresponding URLs are as follows: TCR_priv_/pMHC map EMD-54366 (https://www.ebi.ac.uk/emdb/EMD-54366) and model PDB 9RXM (https://www.rcsb.org/structure/9RXM); TCR_pub_/pMHC map EMD-54262 (https://www.ebi.ac.uk/emdb/EMD-54262) and model PDB 9RU5 (https://www.rcsb.org/structure/9RU5). All data are publicly available for download and visualization from the respective databases.

## Acknowledgements

We would like to thank Proimmune UK provided Peptide-MHC Pentamer. We thank Dr. Jiri Kratochvil for help with the MP analysis. We acknowledge Diamond Light Source for access and support of the cryo-EM facilities at the UK national electron Bio-Imaging Centre (eBIC) (proposal NT29182). Computation was performed at the Oxford Biomedical Research Computing (BMRC) facility supported by the Wellcome Trust Core Award Grant Number 203141/Z/16/Z with additional support from the NIHR Oxford BRC. This work was supported by the Chinese Academy of Medical Sciences (CAMS) Innovation Fund for Medical Science (CIFMS), China (grant number: 2024-I2M-2-001-1) and UK Medical Research Council (MRC) grant MR/Y015347/1 and IMMPROVE - MR/Y004450/1; This work was supported by the National Institutes of Health (U54 AI170791, R21 AI184080), the UK Wellcome Discovery Award (311427/Z/24/Z), and the ERC AdG grant (101021133).

## Author contributions statement

T.D., R.F., and P.Z. conceived the study and designed the experiments. Y.P., D.D., G.L., and X.Y. conducted the immunological assays and performed TCR sequencing. A.J.M. and J.C.K. established the clinical cohort and collected the samples. E.A. carried out the TCR sequence analysis. R.C. and Y.Y contributed with TCR protein production and QC. J.M., Y.P., and J-L.C. purified the proteins, and C.A. and J.M. designed and performed the biochemical characterization experiments for the complexes. C.A. and N.H. prepared the cryo-EM samples. C.A., N.H. and Y.Z. collected the cryo-EM data. Y.Z. processed the TCR_pub_/pMHC dataset and determined the structure; and N.H., Y.Z, C.A processed the TCR_priv_/pMHC dataset and determined the structure. C.A. built and validated atomic models with help from Y.Z., and E.B. C.A. and P.Z. analysed the structures. C.A., R.F., Y.P., and P.Z. prepared the figures, with contributions from E.B., J.M., and Y.Z. Finally, C.A., R.F., Y.P., T.D., and P.Z. wrote the manuscript with input from all authors.

## References

1 Neefjes, J., Jongsma, M. L., Paul, P. & Bakke, O. Towards a systems understanding of MHC class I and MHC class II antigen presentation. Nat Rev Immunol 11, 823–836 (2011). 10.1038/nri3084

2 Zinkernagel, R. M. & Doherty, P. C. Restriction of in vitro T cell-mediated cytotoxicity in lymphocytic choriomeningitis within a syngeneic or semiallogeneic system. Nature 248, 701–702 (1974). 10.1038/248701a0

3 Davis, M. M. & Bjorkman, P. J. T-cell antigen receptor genes and T-cell recognition. Nature 334, 395–402 (1988). 10.1038/334395a0

4 Burnet, F. M. The concept of immunological surveillance. Prog Exp Tumor Res 13, 1–27 (1970). 10.1159/000386035

5 Prehn, R. T. & Main, J. M. Immunity to methylcholanthrene-induced sarcomas. J Natl Cancer Inst 18, 769–778 (1957).

6 van der Bruggen, P. et al. A gene encoding an antigen recognized by cytolytic T lymphocytes on a human melanoma. Science 254, 1643–1647 (1991). 10.1126/science.1840703

7 Tonegawa, S. Somatic generation of antibody diversity. Nature 302, 575–581 (1983). 10.1038/302575a0

8 Arstila, T. P. et al. A direct estimate of the human alphabeta T cell receptor diversity. Science 286, 958–961 (1999). 10.1126/science.286.5441.958

9 Hogquist, K. A. et al. T cell receptor antagonist peptides induce positive selection. Cell 76, 17–27 (1994). 10.1016/0092-8674(94)90169-4

10 Robins, H. S. et al. Comprehensive assessment of T-cell receptor beta-chain diversity in alphabeta T cells. Blood 114, 4099–4107 (2009). 10.1182/blood-2009-04-217604

11 Murugan, A., Mora, T., Walczak, A. M. & Callan, C. G., Jr. Statistical inference of the generation probability of T-cell receptors from sequence repertoires. Proc Natl Acad Sci U S A 109, 16161–16166 (2012). 10.1073/pnas.1212755109

12 Schmidt, C., Burrows, S. R., Sculley, T. B., Moss, D. J. & Misko, I. S. Nonresponsiveness to an immunodominant Epstein-Barr virus-encoded cytotoxic T-lymphocyte epitope in nuclear antigen 3A: implications for vaccine strategies. Proc Natl Acad Sci U S A 88, 9478–9482 (1991). 10.1073/pnas.88.21.9478

13 Argaet, V. P. et al. Dominant selection of an invariant T cell antigen receptor in response to persistent infection by Epstein-Barr virus. J Exp Med 180, 2335–2340 (1994). 10.1084/jem.180.6.2335

14 Dong, T. et al. HIV-specific cytotoxic T cells from long-term survivors select a unique T cell receptor. J Exp Med 200, 1547–1557 (2004). 10.1084/jem.20032044

15 Peng, Y. et al. An immunodominant NP(105-113)-B*07:02 cytotoxic T cell response controls viral replication and is associated with less severe COVID-19 disease. Nat Immunol 23, 50–61 (2022). 10.1038/s41590-021-01084-z

16 Lineburg, K. E. et al. CD8(+) T cells specific for an immunodominant SARS-CoV-2 nucleocapsid epitope cross-react with selective seasonal coronaviruses. Immunity 54, 1055–1065 e1055 (2021). 10.1016/j.immuni.2021.04.006

17 Nesterenko, P. A. et al. HLA-A(*)02:01 restricted T cell receptors against the highly conserved SARS-CoV-2 polymerase cross-react with human coronaviruses. Cell Rep 37, 110167 (2021). 10.1016/j.celrep.2021.110167

18 Augusto, D. G. et al. A common allele of HLA is associated with asymptomatic SARS-CoV-2 infection. Nature 620, 128–136 (2023). 10.1038/s41586-023-06331-x

19 Shomuradova, A. S. et al. SARS-CoV-2 Epitopes Are Recognized by a Public and Diverse Repertoire of Human T Cell Receptors. Immunity 53, 1245–1257 e1245 (2020). 10.1016/j.immuni.2020.11.004

20 Wu, D. et al. Structural assessment of HLA-A2-restricted SARS-CoV-2 spike epitopes recognized by public and private T-cell receptors. Nat Commun 13, 19 (2022). 10.1038/s41467-021-27669-8

21 Liu, G. et al. Long-persisting SARS-CoV-2 spike-specific CD4(+) T cells associated with mild disease and increased cytotoxicity post COVID-19. Nat Commun 16, 8743 (2025). 10.1038/s41467-025-63711-9

22 Nguyen, T. H. O. et al. CD8(+) T cells specific for an immunodominant SARS-CoV-2 nucleocapsid epitope display high naive precursor frequency and TCR promiscuity. Immunity 54, 1066–1082 e1065 (2021). 10.1016/j.immuni.2021.04.009

23 Hong, J. et al. A common TCR V-D-J sequence in V beta 13.1 T cells recognizing an immunodominant peptide of myelin basic protein in multiple sclerosis. J Immunol 163, 3530–3538 (1999).

24 Zeng, W., Maciejewski, J. P., Chen, G. & Young, N. S. Limited heterogeneity of T cell receptor BV usage in aplastic anemia. J Clin Invest 108, 765–773 (2001). 10.1172/jci12687

25 Bentzen, A. K. et al. Large-scale detection of antigen-specific T cells using peptide-MHC-I multimers labeled with DNA barcodes. Nat Biotechnol 34, 1037–1045 (2016). 10.1038/nbt.3662

26 Garcia, K. C. et al. An alphabeta T cell receptor structure at 2.5 A and its orientation in the TCR-MHC complex. Science 274, 209–219 (1996). 10.1126/science.274.5285.209

27 Garcia, K. C. et al. Structural basis of plasticity in T cell receptor recognition of a self peptide-MHC antigen. Science 279, 1166–1172 (1998). 10.1126/science.279.5354.1166

28 Rudolph, M. G., Stanfield, R. L. & Wilson, I. A. How TCRs bind MHCs, peptides, and coreceptors. Annu Rev Immunol 24, 419–466 (2006). 10.1146/annurev.immunol.23.021704.115658

29 Garboczi, D. N. et al. Structure of the complex between human T-cell receptor, viral peptide and HLA-A2. Nature 384, 134–141 (1996). 10.1038/384134a0

30 Stewart-Jones, G. B., McMichael, A. J., Bell, J. I., Stuart, D. I. & Jones, E. Y. A structural basis for immunodominant human T cell receptor recognition. Nat Immunol 4, 657–663 (2003). 10.1038/ni942

31 Gray, G. I. et al. The Evolving T Cell Receptor Recognition Code: The Rules Are More Like Guidelines. Immunol Rev 329, e13439 (2025). 10.1111/imr.13439

32 Lin, V. et al. TCR3d 2.0: expanding the T cell receptor structure database with new structures, tools and interactions. Nucleic Acids Res 53, D604–D608 (2025). 10.1093/nar/gkae840

33 Garcia, K. C., Adams, J. J., Feng, D. & Ely, L. K. The molecular basis of TCR germline bias for MHC is surprisingly simple. Nat Immunol 10, 143–147 (2009). 10.1038/ni.f.219

34 Feng, D., Bond, C. J., Ely, L. K., Maynard, J. & Garcia, K. C. Structural evidence for a germline-encoded T cell receptor-major histocompatibility complex interaction ’codon’. Nat Immunol 8, 975–983 (2007). 10.1038/ni1502

35 Ferretti, A. P. et al. Unbiased Screens Show CD8(+) T Cells of COVID-19 Patients Recognize Shared Epitopes in SARS-CoV-2 that Largely Reside outside the Spike Protein. Immunity 53, 1095–1107 e1093 (2020). 10.1016/j.immuni.2020.10.006

36 Rha, M. S. et al. PD-1-Expressing SARS-CoV-2-Specific CD8(+) T Cells Are Not Exhausted, but Functional in Patients with COVID-19. Immunity 54, 44–52 e43 (2021). 10.1016/j.immuni.2020.12.002

37 Tarke, A. et al. Negligible impact of SARS-CoV-2 variants on CD4 (+) and CD8 (+) T cell reactivity in COVID-19 exposed donors and vaccinees. bioRxiv (2021). 10.1101/2021.02.27.433180

38 Ren, Y. et al. The ORF3a protein of SARS-CoV-2 induces apoptosis in cells. Cell Mol Immunol 17, 881–883 (2020). 10.1038/s41423-020-0485-9

39 Peng, Y. et al. Broad and strong memory CD4(+) and CD8(+) T cells induced by SARS-CoV-2 in UK convalescent individuals following COVID-19. Nat Immunol 21, 1336–1345 (2020). 10.1038/s41590-020-0782-6

40 Kared, H. et al. SARS-CoV-2-specific CD8+ T cell responses in convalescent COVID-19 individuals. J Clin Invest 131 (2021). 10.1172/JCI145476

41 Schulien, I. et al. Characterization of pre-existing and induced SARS-CoV-2-specific CD8(+) T cells. Nat Med 27, 78–85 (2021). 10.1038/s41591-020-01143-2

42 Adamo, S. et al. Signature of long-lived memory CD8(+) T cells in acute SARS-CoV-2 infection. Nature 602, 148–155 (2022). 10.1038/s41586-021-04280-x

43 Gowthaman, R. & Pierce, B. G. TCR3d: The T cell receptor structural repertoire database. Bioinformatics 35, 5323–5325 (2019). 10.1093/bioinformatics/btz517

44 Zhang, Y. & Skolnick, J. TM-align: a protein structure alignment algorithm based on the TM-score. Nucleic Acids Res 33, 2302–2309 (2005). 10.1093/nar/gki524

45 Garcia, K. C., Teyton, L. & Wilson, I. A. Structural basis of T cell recognition. Annu Rev Immunol 17, 369–397 (1999). 10.1146/annurev.immunol.17.1.369

46 Linette, G. P. et al. Cardiovascular toxicity and titin cross-reactivity of affinity-enhanced T cells in myeloma and melanoma. Blood 122, 863–871 (2013). 10.1182/blood-2013-03-490565

47 Petrova, G., Ferrante, A. & Gorski, J. Cross-reactivity of T cells and its role in the immune system. Crit Rev Immunol 32, 349–372 (2012). 10.1615/critrevimmunol.v32.i4.50

48 Dong, D. et al. Structural basis of assembly of the human T cell receptor-CD3 complex. Nature 573, 546–552 (2019). 10.1038/s41586-019-1537-0

49 Susac, L. et al. Structure of a fully assembled tumor-specific T cell receptor ligated by pMHC. Cell 185, 3201–3213 e3219 (2022). 10.1016/j.cell.2022.07.010

50 Xin, W. et al. Structures of human gammadelta T cell receptor-CD3 complex. Nature 630, 222–229 (2024). 10.1038/s41586-024-07439-4

51 Saotome, K. et al. Structural analysis of cancer-relevant TCR-CD3 and peptide-MHC complexes by cryoEM. Nat Commun 14, 2401 (2023). 10.1038/s41467-023-37532-7

52 Sewell, A. K. Why must T cells be cross-reactive? Nat Rev Immunol 12, 669–677 (2012). 10.1038/nri3279

53 Emerson, R. O. et al. Immunosequencing identifies signatures of cytomegalovirus exposure history and HLA-mediated effects on the T cell repertoire. Nat Genet 49, 659–665 (2017). 10.1038/ng.3822

54 Milighetti, M. et al. Large clones of pre-existing T cells drive early immunity against SARS-COV-2 and LCMV infection. iScience 26, 106937 (2023). 10.1016/j.isci.2023.106937

55 Picelli, S. et al. Full-length RNA-seq from single cells using Smart-seq2. Nat Protoc 9, 171–181 (2014). 10.1038/nprot.2014.006

56 Liao, Y., Smyth, G. K. & Shi, W. featureCounts: an efficient general purpose program for assigning sequence reads to genomic features. Bioinformatics 30, 923–930 (2014). 10.1093/bioinformatics/btt656

57 Stuart, T. et al. Comprehensive Integration of Single-Cell Data. Cell 177, 1888–1902 e1821 (2019). 10.1016/j.cell.2019.05.031

58 Dobin, A. et al. STAR: ultrafast universal RNA-seq aligner. Bioinformatics 29, 15–21 (2013). 10.1093/bioinformatics/bts635

59 Bolotin, D. A. et al. Antigen receptor repertoire profiling from RNA-seq data. Nat Biotechnol 35, 908–911 (2017). 10.1038/nbt.3979

60 Bolotin, D. A. et al. MiXCR: software for comprehensive adaptive immunity profiling. Nat Methods 12, 380–381 (2015). 10.1038/nmeth.3364

61 Abd Hamid, M., et al. Self-Maintaining CD103(+) Cancer-Specific T Cells Are Highly Energetic with Rapid Cytotoxic and Effector Responses. Cancer Immunol Res 8, 203–216 (2020). 10.1158/2326-6066.Cir-19-0554

62 Brunson, J. C. ggalluvial: Layered Grammar for Alluvial Plots. J Open Source Softw 5 (2020). 10.21105/joss.02017

63 Garboczi, D. N., Hung, D. T. & Wiley, D. C. HLA-A2-peptide complexes: refolding and crystallization of molecules expressed in Escherichia coli and complexed with single antigenic peptides. Proc Natl Acad Sci U S A 89, 3429–3433 (1992). 10.1073/pnas.89.8.3429

64 Punjani, A., Rubinstein, J. L., Fleet, D. J. & Brubaker, M. A. cryoSPARC: algorithms for rapid unsupervised cryo-EM structure determination. Nat Methods 14, 290–296 (2017). 10.1038/nmeth.4169

65 Abramson, J. et al. Addendum: Accurate structure prediction of biomolecular interactions with AlphaFold 3. Nature 636, E4 (2024). 10.1038/s41586-024-08416-7

66 Emsley, P., Lohkamp, B., Scott, W. G. & Cowtan, K. Features and development of Coot. Acta Crystallogr D Biol Crystallogr 66, 486–501 (2010). 10.1107/S0907444910007493

67 Afonine, P. V. et al. Real-space refinement in PHENIX for cryo-EM and crystallography. Acta Crystallogr D Struct Biol 74, 531–544 (2018). 10.1107/S2059798318006551

68 Pettersen, E. F. et al. UCSF ChimeraX: Structure visualization for researchers, educators, and developers. Protein Sci 30, 70–82 (2021). 10.1002/pro.3943

69 Edgar, R. C. MUSCLE: multiple sequence alignment with high accuracy and high throughput. Nucleic Acids Res 32, 1792–1797 (2004). 10.1093/nar/gkh340

70 Robert, X. & Gouet, P. Deciphering key features in protein structures with the new ENDscript server. Nucleic Acids Res 42, W320–324 (2014). 10.1093/nar/gku316

