## supplemental tables and figures for "Uncovering structural determinants of peptide recognition by public and private T-cell receptors"

### Supplementary tables and figures

**Supplementary Table 1**. T cell response against epitope HLA-A*01:01 ORF3a$_{(207-215)}$ measured by *ex vivo* IFN-γ ELISpot. SFU: Spot Forming Unit from 18 HLA-A*01:01 positive individuals.

| Participant ID | HLA-A*01:01 | Responder | SFU/10^6PBMCs |
|---|---|---|---|
| C-COV19-001 | No | | |
| C-COV19-002 | No | | |
| C-COV19-003 | No | | |
| C-COV19-004 | No | | |
| C-COV19-005 | Yes | Yes | 25 |
| C-COV19-006 | Yes | Yes | 30 |
| C-COV19-007 | Yes | Yes | 25 |
| C-COV19-008 | No | | |
| C-COV19-009 | No | | |
| C-COV19-010 | No | | |
| C-COV19-011 | Yes | No | 0 |
| C-COV19-012 | No | | |
| C-COV19-014 | Yes | Yes | 15 |
| C-COV19-018 | No | | |
| C-COV19-020 | No | | |
| C-COV19-021 | No | | |
| C-COV19-022 | Yes | Yes | 50 |
| C-COV19-023 | No | | |
| C-COV19-024 | No | | |
| C-COV19-025 | No | | |
| C-COV19-026 | No | | |
| C-COV19-027 | No | | |
| C-COV19-028 | No | | |
| C-COV19-029 | No | | |
| C-COV19-030 | No | | |
| C-COV19-031 | No | | |
| C-COV19-032 | No | | |
| C-COV19-033 | No | | |
| C-COV19-034 | Yes | Yes | 35 |
| C-COV19-035 | No | | |
| C-COV19-036 | Yes | Yes | 120 |
| C-COV19-037 | Yes | Yes | 115 |
| C-COV19-038 | No | | |
| C-COV19-039 | Yes | Yes | 60 |
| C-COV19-040 | Yes | Yes | 90 |
| C-COV19-041 | Yes | Yes | 125 |
| C-COV19-042 | No | | |
| C-COV19-043 | No | | |
| C-COV19-044 | No | | |
| C-COV19-045 | Yes | Yes | 125 |
| C-COV19-046 | No | | |
| C-COV19-047 | No | | |
| C-COV19-048 | No | | |
| C-COV19-050 | No | | |
| C-COV19-051 | Yes | Yes | 40 |
| C-COV19-054 | Yes | NO | 0 |
| C-COV19-055 | No | | |
| C-COV19-056 | Yes | Yes | 35 |
| C-COV19-059 | Yes | Yes | 110 |
| C-COV19-060 | No | | |
| C-COV19-061 | No | | |
| C-COV19-062 | Yes | Yes | 40 |

**Supplementary Table 2**. Clonotype and CDR3 amino acid sequences of $TCR_{pub}$ and $TCR_{priv}$

| ID | TRBV | CDR3_beta | TRAV | CDR3_alpha | TRAJ | Va_ public | Paired_ public |
|---|---|---|---|---|---|---|---|
| **$TCR_{pub}$** | TRBV5-1 | CASSLAGDLGTEAFF | TRAV3 | CAVRVVTSGGSYIPTF | TRAJ6 | Public | Public |
| **$TCR_{priv}$** | TRBV5-1 | CASSLWTGVGTEAFF | TRAV25 | CASDREGGSEKLVF | TRAJ57 | Private | Private |

**Supplementary Table 3**. Cryo-EM data collection and structure determination of TCR/pMHC complexes

| **Data collection and processing** | | |
|---|---|---|
| Complex | **$TCR_{pub}$/pMHC**<br>EMDB-54262<br>PDB_ID: 9RU5 | **$TCR_{priv}$/pMHC**<br>EMDB-54336<br>PDB_ID: 9RXM |
| Microscope | Titan Krios | |
| Voltage | 300 kV | |
| Electron dose | 50 e-/Å$^2$ | |
| Detector | Gatan K3 | Falcon<br>4i SelectrisX |
| Defocus Range | -0.8 to -2.5 μm | |
| Pixel Size | 0.825 | 0.7303 |
| Micrograph numbers | 12657 | 10222 |
| Particle numbers | 5,535,316 | 3,903,441 |
| Symmetry | C1 | C1 |
| Map resolution (Å) | 3.4 | 3.37 |
| FSC threshold | 0.143 | 0.143 |
| Local resolution range (Å) | 2.92-11.21 | 2.5-9.67 |
| **Refinement** | | |
| Initial Model used (PDB code) | AlphaFold3 Prediction | |
| FSC threshold | 0.143 | 0.143 |
| Non-hydrogen atoms | 4825 | 6063 |
| Residues | 605 | 752 |
| Bond length(Å) | 0.003 | 0.003 |
| Bond angles(°) | 0.699 | 0.642 |
| MolProbity score | 2.19 | 2.19 |
| Clashscore | 17.33 | 17.33 |
| Rotamers Outliers (%) | 0 | 0 |
| Ramachandran | | |
| Favored(%) | 92.84 | 88.41 |
| Allowed(%) | 6.98 | 11.45 |
| Outliers(%) | 0.17 | 0.14 |

**Supplementary Table 4 .** Structural comparison and RMSD analysis of $TCR_{pub}$ and $TCR_{priv}$ complexes. This table contains the RMSD values of the PDB structures used in Figure 6c. HLA alleles highlighted in white correspond to the same subtype, while the others (in grey) represent different subtypes.

| $TCR_{pub}$ Chains | Target PDB | Target Chains | CA Atoms | RMSD (Å) | HLA type | peptide |
|---|---|---|---|---|---|---|
| A&B | 5E6I | P&Q | 233 | 9.117 | HLA-A*02:01 | GILGFVFTL |
| A&B | 8WUL | C&D | 237 | 9.632 | HLA-A*11:01 | VVGAVGVGK |
| A&B | 6UON | G&H | 235 | 10.192 | HLA-C*08:02 | GADGVGKSAL |
| A&B | 8WTE | A&B | 234 | 11.021 | HLA-A*11:01 | VVGAVGVGK |
| A&B | 7PHR | A&B | 236 | 12.202 | HLA-A*02:01 | YLEPGPVTV |
| A&B | 8I5D | A&B | 235 | 12.657 | HLA-A*11:01 | VVGAVGVGK |
| A&B | 4MVB | C&D | 236 | 13.854 | H-2 $L^d$ (mouse) | QPAEGGFQL |
| A&B | 6BJ2 | D&E | 237 | 13.878 | HLA-B*35:01 | IPLTEEAEL |
| A&B | 3RGV | A&B | 236 | 13.99 | H-2 $K^b$ (mouse) | WIYVYRPMGCGGS |
| A&B | 6AVF | A&B | 236 | 14.142 | HLA-B*07:02 | APRGPHGGAASGL |
| A&B | 4MS8 | C&D | 236 | 14.235 | H-2 $L^d$ (mouse) | SPAEAGFFL |
| A&B | 7N1F | D&E | 236 | 14.492 | HLA-A*02:01 | YLQPRTFLL |
| A&B | 8GVB | A&B | 228 | 14.598 | HLA-A*24:02 | RYPLTFGW |
| A&B | 8GVG | A&B | 235 | 14.918 | HLA-A*24:02 | RFPLTFGW |
| A&B | 8GVI | A&B | 235 | 15.038 | HLA-A*24:02 | RYPLTFGW |
| A&B | 8ES9 | A&B | 236 | 15.136 | HLA-A*02:01 | GVYDGREHTV |
| A&B | 5D2N | C&F | 237 | 16.405 | HLA-A*02:01 | NLVPMVATV |
| A&B | 6AVG | B&E | 235 | 16.468 | HLA-B*07:02 | APRGPHGGAASGL |
| A&B | 7N2S | D&F | 236 | 16.689 | HLA-B*27 | TRLALIAPK |
| A&B | 2E7L | B&C | 119 | 16.783 | H-2 $L^d$ (mouse) | QLSPFPFDL |
| A&B | 3TPU | A&B | 237 | 16.905 | H-2 $L^d$ (mouse) | FLSPFWFDI |
| A&B | 8D5Q | A&B | 237 | 16.917 | H-2 $L^d$ (mouse) | HPGSVNEFDF |
| A&B | 7N2N | D&F | 236 | 17.058 | HLA-B*27 | TRLALIAPK |
| A&B | 7N2R | D&F | 236 | 17.598 | HLA-B*27. | TRLALIAPK |
| A&B | 4NHU | C&D | 236 | 17.607 | H-2 $L^d$ (mouse) | GGGAPWNPAMMI |
| A&B | 7N2P | D&F | 236 | 17.919 | HLA-B*27 | GQVMVVAPR |
| A&B | 5YXN | A&B | 236 | 18.018 | HLA-A*02:01 | KLVALGINAV |
| A&B | 7N2Q | D&F | 236 | 18.521 | HLA-B*27 | LRVMMLAPF |
| A&B | 2CKB | A&B | 237 | 18.713 | H-2 $L^d$ (mouse) | EQYKFYSV |
| A&B | 1G6R | A&B | 237 | 19.231 | H-2 $K^b$ (mouse) | SIYRYYGL |
| A&B | 1NAM | A&B | 116 | 19.534 | H-2 $K^b$ (mouse) | RGYVYQGL |
| A&B | 2OL3 | A&B | 118 | 19.575 | H-2 $K^b$ (mouse) | SQYYYNSL |
| A&B | 1MWA | A&B | 237 | 19.582 | H-2 $K^b$ (mouse) | EQYKFYSV |
| A&B | 8RRO | A&B | 235 | 19.604 | HLA-A*03:01 | VVVGAVGVGK |
| A&B | 7RE7 | H&L | 217 | 20.275 | HLA-A*02:01 | FMNKFIYEI |
| A&B | 7N2O | D&F | 236 | 20.444 | HLA-B*27 | LRVMMLAPF |
| A&B | 4MXQ | C&D | 237 | 21.055 | H-2 $L^d$ (mouse) | SPAPRPLDL |
| A&B | 5TEZ | I&J | 235 | 22.154 | HLA-A*02:01 | GILGFVFTL |
| A&B | 4N5E | C&D | 237 | 22.341 | H-2 $L^d$ (mouse) | VPYMAEFGM |
| A&B | 3MV9 | D&E | 226 | 22.906 | HLA-B*35:01 | HPVGEADYFEY |
| A&B | 7RTR | D&E | 225 | 23.265 | HLA-A*02:01 | YLQPRTFLL |
| A&B | 7N6E | I&J | 225 | 25.525 | HLA-A*02:01 | YLQPRTFLL |
| A&B | 7JWJ | E&D | 224 | 26.619 | H-2 $L^d$ (mouse) | ASNENMETM |
| A&B | 7STF | H&L | 221 | 27.832 | HLA-A*03:01. | VVVGAVGVGK |
| A&B | 8ENH | D&E | 228 | 28.754 | HLA-B*35:01 | LPFEKSTIM |
| A&B | 9C3E | A&B | 217 | 30.133 | HLA-A*02:01 | SLLMWITQV |
| A&B | 8FJA | E&D | 221 | 32.795 | HLA-A*02:01 | GVYDGREHTV |
| A&B | 3CII | H&G | 170 | 35.608 | HLA-E | VMAPRTLFL |
| A&B | 4N8V | G | 190 | 36.473 | HLA-A*11:01 | MLIYSMWGK |
| A&B | 8T7R | I&J | 225 | 56.115 | HLA-A*01:01 | VTEHDTLLY |

| $TCR_{priv}$ Chains | Target PDB | Target Chains | CA Atoms | RMSD (Å) | HLA type | peptide |
|---|---|---|---|---|---|---|
| A&B | 8WTE | C&D | 221 | 8.602 | HLA-A*11:01 | VVGAVGVGK |
| A&B | 8WUL | C&D | 221 | 8.805 | HLA-A*11:01 | VVGAVGVGK |
| A&B | 5E6I | G&H | 219 | 9.483 | HLA-A*02:01 | GILGFVFTL |
| A&B | 7PHR | A&B | 220 | 9.566 | HLA-A*02:01 | YLEPGPVTV |
| A&B | 6UON | G&H | 214 | 9.892 | HLA-C*08:02 | GADGVGKSAL |
| A&B | 8I5D | A&B | 219 | 10.404 | HLA-A*11:01 | VVGAVGVGK |
| A&B | 8FJA | A&B | 220 | 11.12 | HLA-A*02:01 | GVYDGREHTV |
| A&B | 3RGV | A&B | 220 | 11.137 | H-2 $K^b$ (mouse) | WIYVYRPMGCGGS |
| A&B | 7N1F | D&E | 220 | 11.943 | HLA-A*02:01 | YLQPRTFLL |
| A&B | 6BJ2 | D&E | 221 | 12.25 | HLA-B*35:01 | IPLTEEAEL |
| A&B | 4MVB | C&D | 220 | 12.386 | H-2 $L^d$ (mouse) | QPAEGGFQL |
| A&B | 4MS8 | C&D | 220 | 13.426 | H-2 $L^d$ (mouse) | SPAEAGFFL |
| A&B | 8GVB | A&B | 212 | 13.444 | HLA-A*24:02 | RYPLTFGW |
| A&B | 8GVI | A&B | 219 | 13.571 | HLA-A*24:02 | RYPLTFGW |
| A&B | 8GVG | A&B | 219 | 13.634 | HLA-A*24:02 | RFPLTFGW |
| A&B | 3MV7 | D&E | 115 | 13.872 | HLA-B*35:01 | HPVGEADYFEY |
| A&B | 5TEZ | D&F | 220 | 14.043 | HLA-A*02:01 | GILGFVFTL |
| A&B | 3TPU | A&B | 221 | 14.078 | H-2 $L^d$ (mouse) | FLSPFWFDI |
| A&B | 7N2O | D&F | 220 | 14.592 | HLA-B*27 | LRVMMLAPF |
| A&B | 2E7L | B&C | 119 | 14.621 | H-2 $L^d$ (mouse) | QLSPFPFDL |
| A&B | 8D5Q | A&B | 221 | 14.743 | H-2 $L^d$ (mouse) | HPGSVNEFDF |
| A&B | 5YXU | A&B | 221 | 14.806 | HLA-A*02:01 | KLVALGINAV |
| A&B | 5YXN | A&B | 220 | 14.822 | HLA-A*02:01 | KLVALGINAV |
| A&B | 5D2N | C&F | 221 | 14.978 | HLA-A*02:01 | NLVPMVATV |
| A&B | 6AVF | B&E | 219 | 14.979 | HLA-B*07:02 | APRGPHGGAASGL |
| A&B | 7N2R | D&F | 220 | 15.124 | HLA-B*27 | TRLALIAPK |
| A&B | 4N8V | U&J | 219 | 15.498 | HLA-A*11:01 | MLIYSMWGK |
| A&B | 4EUP | D&F | 220 | 15.684 | HLA-A*02:01 | ALGIGILTV |
| A&B | 4NHU | C&D | 220 | 15.772 | H-2 $L^d$ (mouse) | GGGAPWNPAMMI |
| A&B | 7N2Q | D&F | 220 | 16.1 | HLA-B*27 | LRVMMLAPF |
| A&B | 2CKB | A&B | 221 | 16.583 | H-2 $K^b$ (mouse) | EQYKFYSV |
| A&B | 1NAM | A&B | 116 | 16.602 | H-2 $K^b$ (mouse) | RGYVYQGL |
| A&B | 8RRO | F&G | 220 | 16.814 | HLA-A*03:01 | VVVGAVGVGK |
| A&B | 8ES9 | A&B | 220 | 16.962 | HLA-A*02:01 | GVYDGREHTV |
| A&B | 1G6R | A&B | 221 | 17.059 | H-2 $K^b$ (mouse) | SIYRYYGL |
| A&B | 2OL3 | A&B | 118 | 17.327 | H-2 $K^b$ (mouse) | SQYYYNSL |
| A&B | 1MWA | A&B | 221 | 17.439 | H-2 $K^b$ (mouse) | EQYKFYSV |
| A&B | 6AVG | H&L | 204 | 17.637 | HLA-B*07:02 | APRGPHGGAASGL |
| A&B | 7N2S | D&F | 220 | 18.41 | HLA-B*27 | TRLALIAPK |
| A&B | 4MXQ | C&D | 221 | 19.72 | H-2 $L^d$ (mouse) | SPAPRPLDL |
| A&B | 4PRH | I&J | 219 | 19.885 | HLA-B*35:01 | HPVGDADYFEY |
| A&B | 4N5E | C&D | 221 | 20.866 | H-2 $L^d$ (mouse) | VPYMAEFGM |
| A&B | 7N2P | D&E | 210 | 21.84 | HLA-B*27 | GQVMVVAPR |
| A&B | 7N6E | D&E | 210 | 22.06 | HLA-A*02:01 | YLQPRTFLL |
| A&B | 5E9D | I&J | 209 | 24.047 | HLA-A*0201 | ELAGIGILTV |
| A&B | 9C3E | A&B | 161 | 25.437 | HLA-A*02:01 | SLLMWITQV |
| A&B | 8T7R | H&L | 208 | 25.86 | HLA-A*01:01 | VTEHDTLLY |
| A&B | 7STF | E&D | 208 | 32.411 | HLA-A*03:01 | VVVGAVGVGK |
| A&B | 7N2N | G | 183 | 34.794 | HLA-B*27 | TRLALIAPK |
| A&B | 7RE7 | R&Q | 210 | 56.516 | HLA-A*02:01 | FMNKFIYEI |

**Supplementary Table 5**. Summary of key TCR–pMHC contacts identified in the cryo-EM structures of $TCR_{pub}$ and $TCR_{priv}$ complexes.

| Complex | TCR Chain | CDR Loop | TCR Residue | Peptide/MHC Residue | Interaction Type |
|---|---|---|---|---|---|
| TCRpub/pMHC | β | CDR2β | Arg53 | Asp4 (peptide) | Hydrogen bond |
| TCRpub/pMHC | β | CDR3β | Ala94 | Tyr5 (peptide) | Hydrophobic |
| TCRpub/pMHC | α | CDR3α | Gly99 | Gln153 (MHC α2 helix) | Hydrogen bond |
| TCRpub/pMHC | α | CDR3α | Val95 | Val156 (MHC α2 helix) | Hydrophobic |
| TCRpub/pMHC | β | CDR2β | Thr51 | Arg63 (MHC α1 helix) | Hydrophobic |
| TCRpub/pMHC | α | CDR1α | Tyr31 | Arg155 (MHC α2 helix) | Hydrogen bond |
| TCRpriv/pMHC | β | CDR2β | Arg53 | Tyr6 (peptide) | Hydrogen bond |
| TCRpriv/pMHC | β | CDR2β | Arg53 | Asp4 (peptide) | Salt bridge |
| TCRpriv/pMHC | β | CDR3β | Trp94 | Leu8 (peptide) | Hydrophobic |
| TCRpriv/pMHC | α | CDR3α | Glu93 | Arg161 (MHC α2 helix) | Hydrogen bond |
| TCRpriv/pMHC | α | CDR2α | Gln47 | Gln153 (MHC α2 helix) | Hydrogen bond |
| TCRpriv/pMHC | β | CDR2β | Asn54 | Arg63 (MHC α1 helix) | Salt bridge |
| TCRpriv/pMHC | α | CDR1α | Thr27 | Val156 (MHC α2 helix) | Hydrophobic |

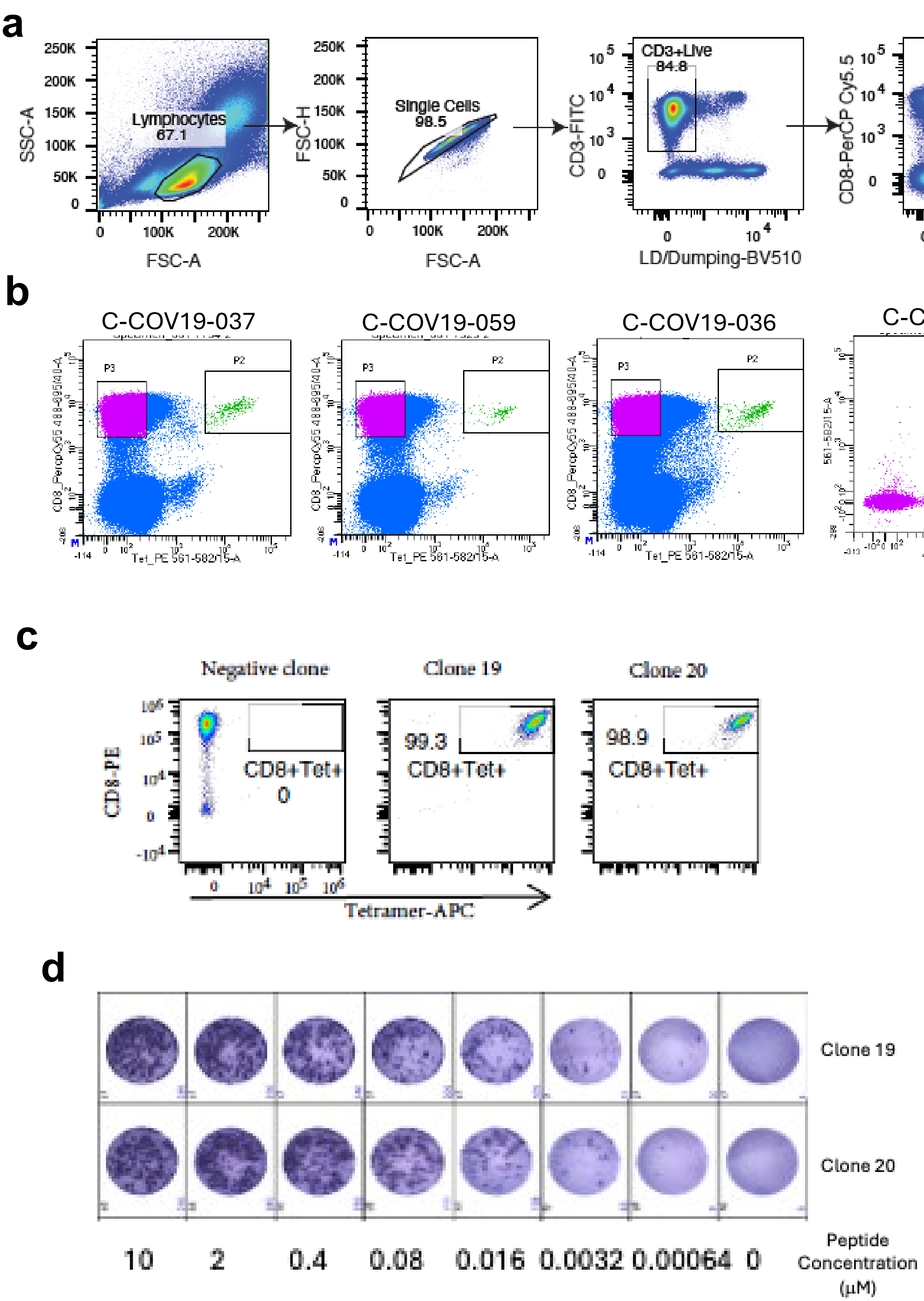

**Supplementary Figure 1.** Flow cytometry data. (**a**) Gating strategy used for flow cytometric sorting of HLA-A01:01/ORF3a$_{(207-215)}$-specific CD8$^{+}$ T cells. (**b**) Representative FACS plots showing the sorted Pentamer$^{+}$ cell populations. (**c**) HLA-A01:01/ORF3a$_{(207-215)}$-specific CD8$^{+}$ T cells were single-cell sorted and expanded *in vitro* as T cell clones. The purity of the clones after expansion was confirmed by staining with peptide–MHC tetramers. (**d**) Representative ELISpot image from the EC$_{50}$ assay. T cell clones were co-cultured with autologous EBV-transformed B cell lines (BCLs) pre-loaded with peptides at fivefold serial dilutions starting from 10 μM.

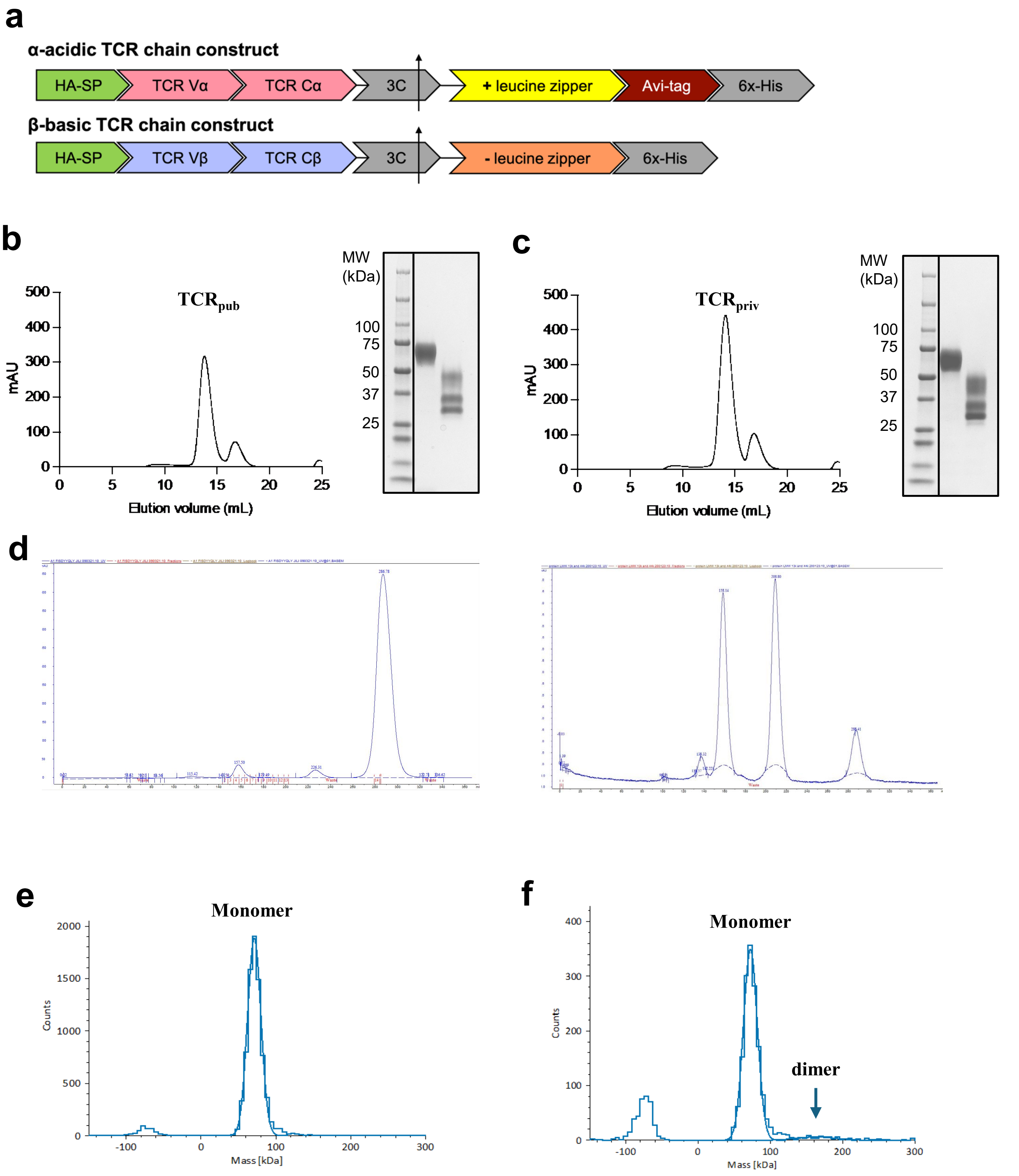

**Supplementary Figure 2.** Expression of soluble ORF3a$_{207\text{-}215}$-A*01:01 TCRs. **(a)** Schematic of a generic soluble TCRαβ DNA construct. **(b-c)** Size-exclusion chromatography profile (left) and SDS-PAGE analysis (right, under reducing and non-reducing conditions) of the 3C-treated soluble ORF3a-reactive TCR$_{pub}$ **(b)** and TCR$_{priv.}$ **(c).** The left peak eluting at approximately 14.0 mL corresponds to the ORF3a TCRs, and the right peak eluting after 16 mL corresponds to the cleaved zippers and tags. **(d)** Gel filtration chromatograms showing the purification of the refolded peptide–MHC complex (top) and the calibration standards (bottom). For calibration, ovalbumin (43 kDa) eluted at 158.14 mL and ribonuclease A (13.7 kDa) at 208.8 mL. The pMHC complex (45 kDa) eluted as a peak at 157.50 mL, whereas free β$_2$M (12 kDa) eluted at 226.31 mL.

a

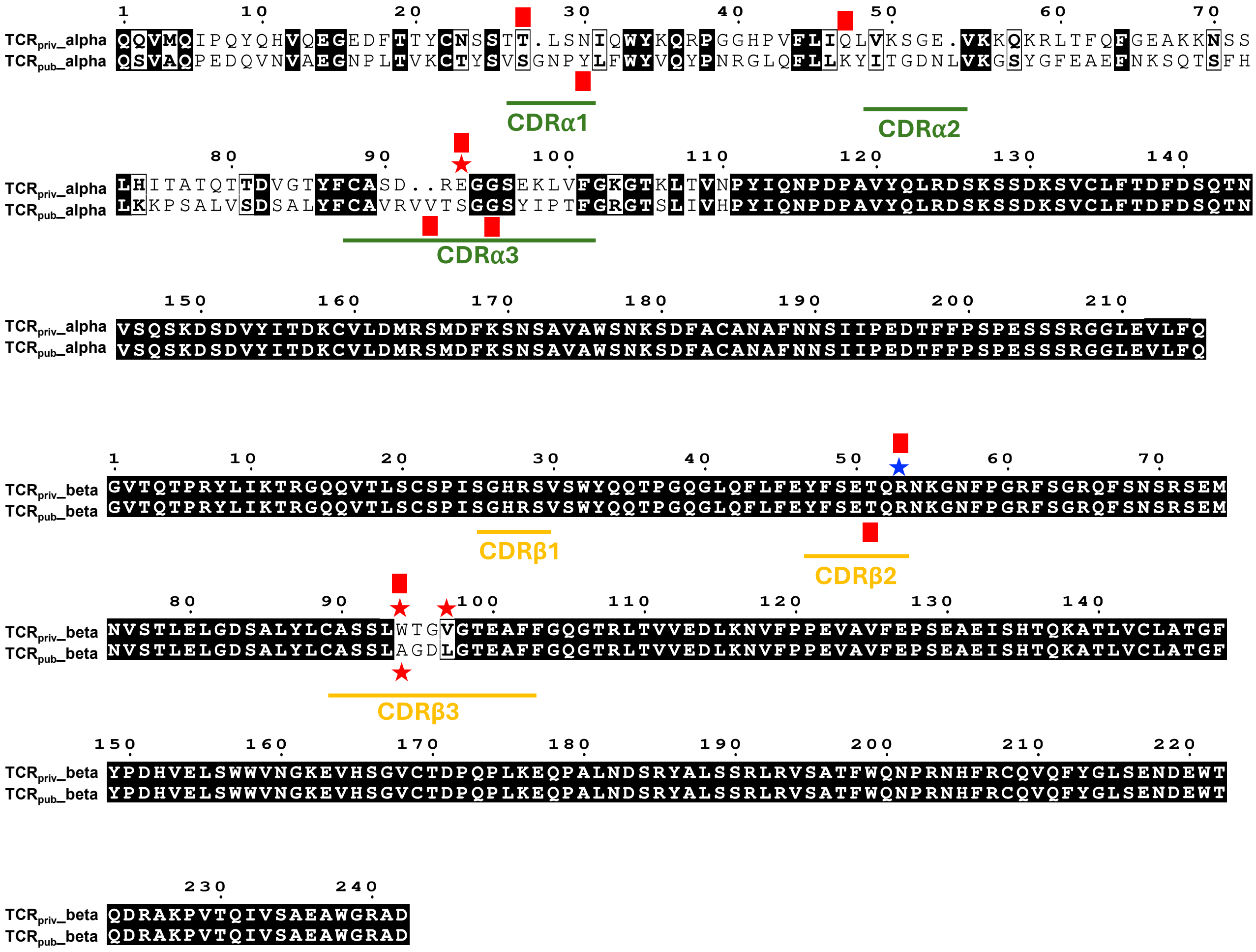

b

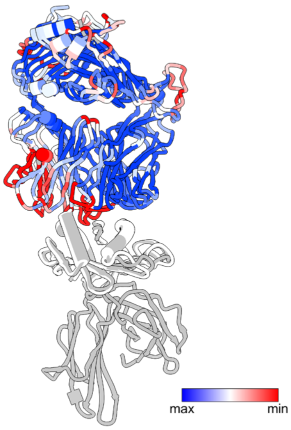

**Supplementary Figure 3.** (**a**) Sequence alignment of the $TCR_{pub}$ and $TCR_{priv}$ α (top) and β (bottom) chains. Key residues involved in peptide interaction are marked with asterisks: red asterisks indicate peptide-contacting residues specific to the α or β chain of $TCR_{pub}$ or $TCR_{priv}$, while blue asterisks denote peptide-contacting residues shared between the α and β chains of both receptors. Residues involved in MHC interaction are boxed: red boxes indicate MHC-contacting residues specific to the α or β chain of $TCR_{priv}$ or $TCR_{priv}$. CDR loops are colored by chain type (green, α; yellow, β). The sequence alignment was performed using MUSCLE and visualized with ESPript 3.0. (**b**) $TCR_{pub}$ aligned with the $TCR_{priv}$, with an RMSD of 1.134 Å. Sequence alignments of TCR-pMHC structures are color-coded based on residue conservation: blue represents highly conserved residues, while red indicates the least conserved.

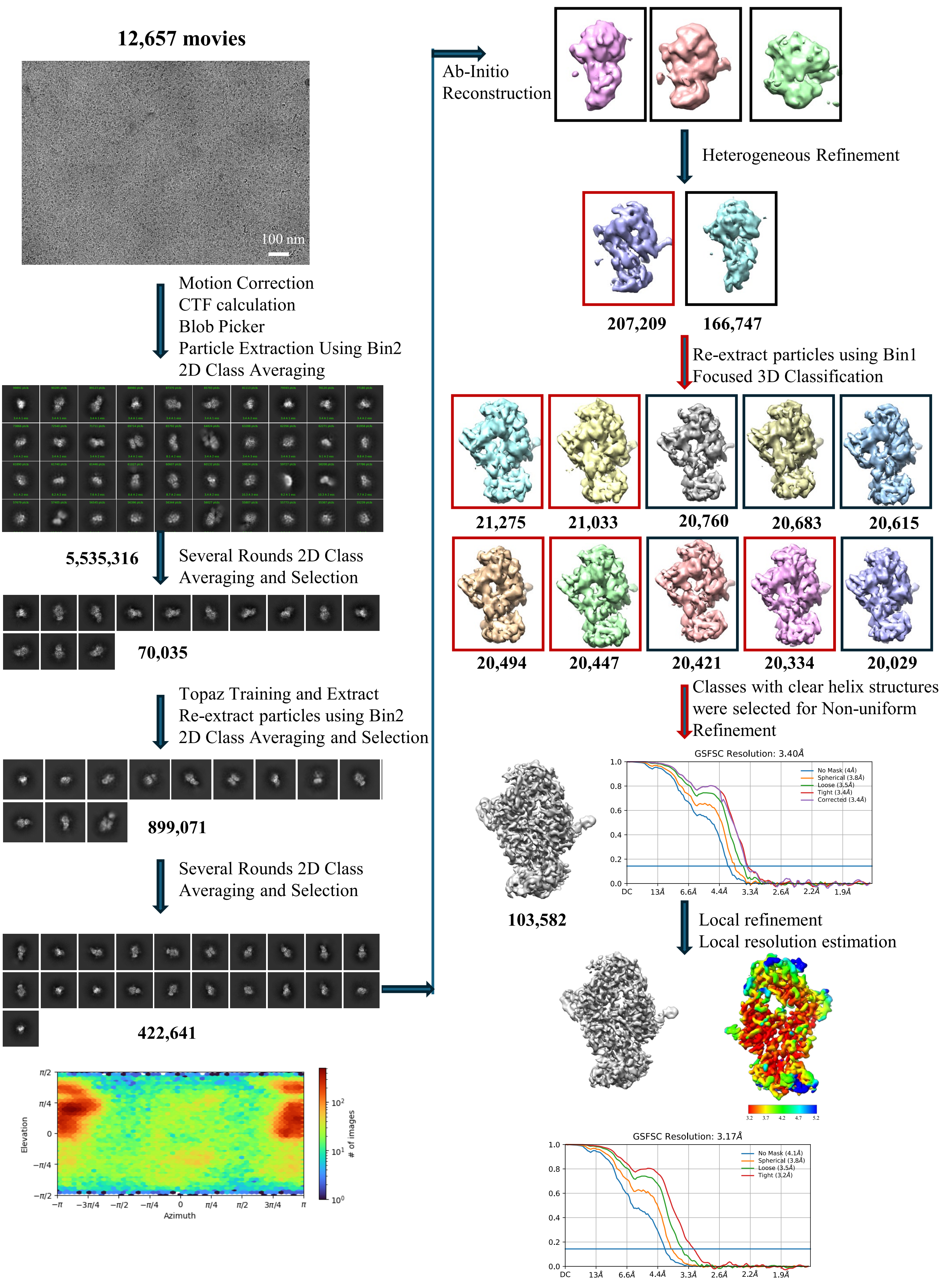

**Supplementary Figure 4.** Single particle cryo-EM data processing workflow for the $TCR_{pub}$/pMHC complex.

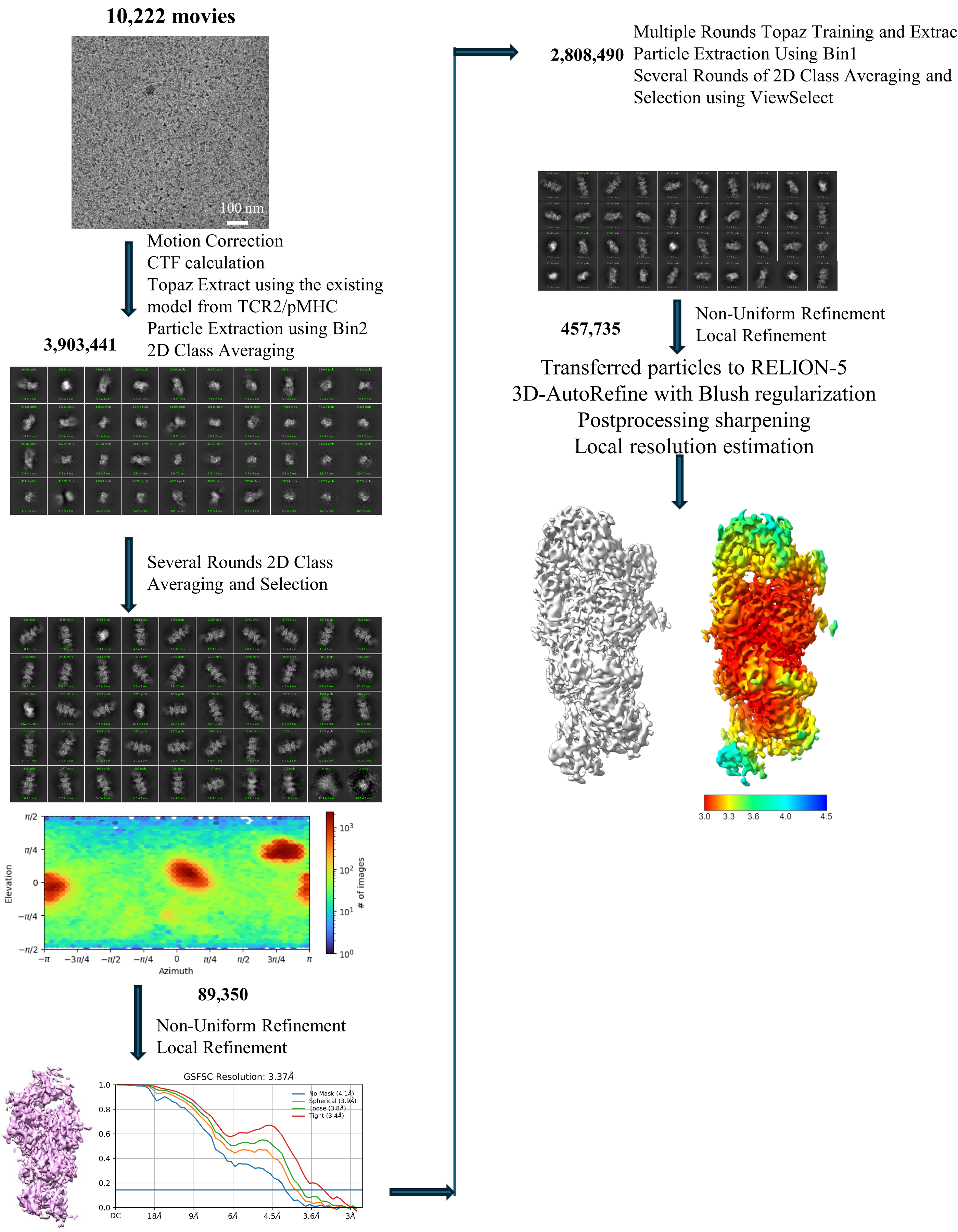

**Supplementary Figure 5.** Single particle cryo-EM data processing workflow for the $TCR_{priv}$/pMHC complex.

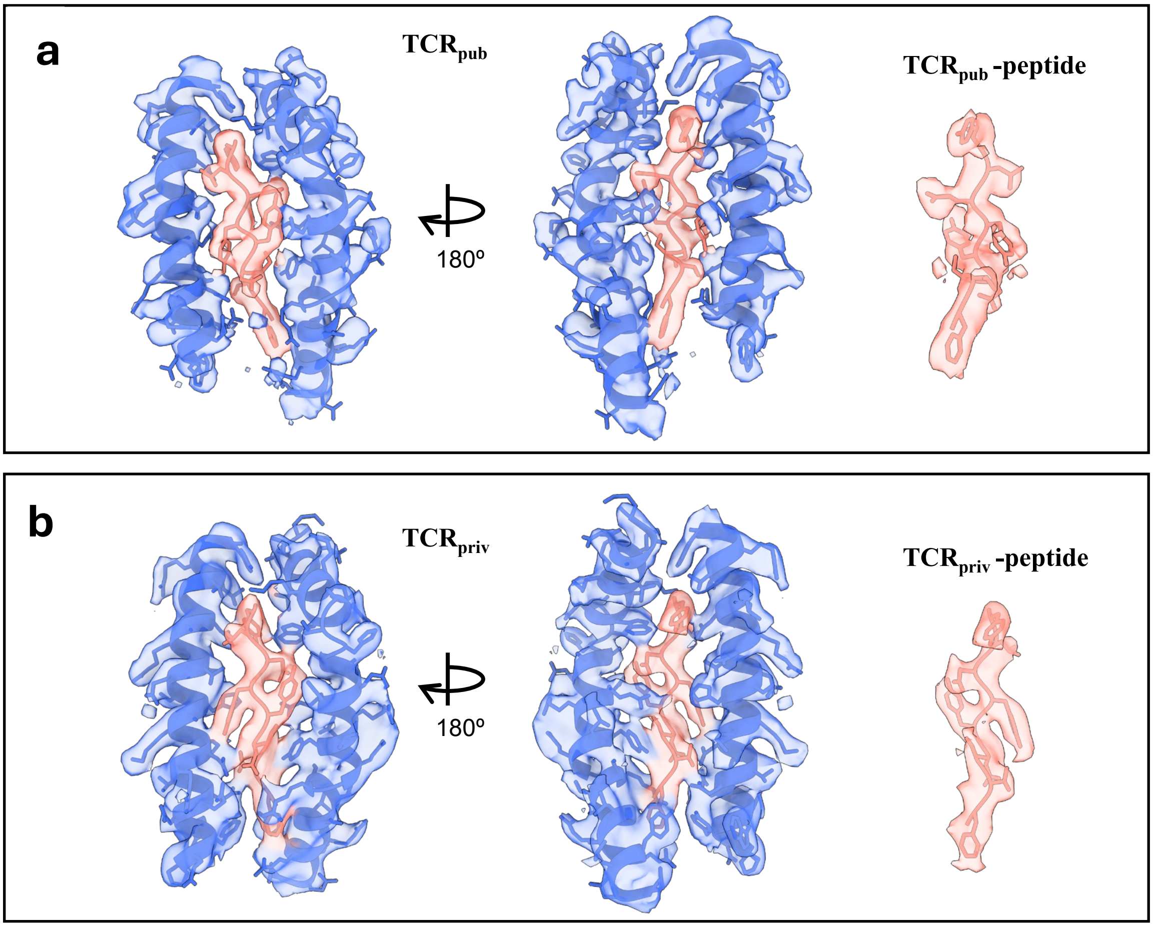

**Supplementary Figure 6. Alternative views of Figures 4a and 4d**. Structural details of (a) the TCR$_{pub}$/pMHC and (b) the TCR$_{priv}$/pMHC complexes, showing the ORF3a(207–215) peptide and MHC interface, with the cryo-EM density map and atomic model superimposed.
